# Episymbiotic Saccharibacteria suppresses epithelial immunoactivation through Type IV pili and TLR2 dependent endocytosis

**DOI:** 10.1101/2025.05.30.656655

**Authors:** Deepak Chouhan, Alex S. Grossman, Kristopher A. Kerns, Kendall S. Stocke, Maya Kim, Pu-Ting Dong, Ajay Kumar, Lei Lei, Richard J. Lamont, Jeffrey S. McLean, Xuesong He, Batbileg Bor

## Abstract

Saccharibacteria are episymbionts that require host-bacteria to grow. They are positively associated with inflammatory diseases within the human microbiome, yet their mechanisms for interacting with the human host and contributing to diseases remain unknown. This study investigated interactions between a Saccharibacterium (*Nanosynbacter lyticus*), its host-bacteria (*Schaalia odontolytica*), and oral epithelial cells. The host-bacteria induced proinflammatory cytokines in epithelial cells, while Saccharibacteria were immune silent. Remarkably, Saccharibacteria dampened cytokine responses to host-bacteria during coinfection. This effect was driven by Saccharibacteria-induced clustering of TLR2 receptors, a process likely facilitated by type IV, ultimately leading to reduced TLR2-mediated cytokine signalling. High resolution imaging showed that Saccharibacteria were endocytosed by oral epithelial cells, and colocalized with endosome markers, eventually trafficking to lysosomes. Moreover, a subset of the Saccharibacteria survive endocytosis long-term, and retains their capability to reinfect host-bacteria, highlighting a mechanism for persistence in the oral microbiome and a vital role in mammalian immune system modulation.

## Introduction

The superphylum Patescibacteria, also known as the Candidate Phyla Radiation or CPR, comprises over 73 clades, representing anywhere between 15–50% of total bacterial diversity^1–5^. Clades Absconditabacteria (SR1), Gracilibacteria (GN02), and Saccharibacteria (TM7) are consistently associated with the mammalian oral cavity and digestive system^2,6,7^. Saccharibacteria are the most extensively studied, due to their widespread presence in both the environment and human microbiome, as well as the availability of culturable strains^8–10^. A total of 7 species and 47 strains have been cultured from the human oral cavity, all having an obligate episymbiotic lifestyle, wherein they grow on the surface of Actinobacteria hosts (e.g., Schaalia, *Actinomyces*)^6,11–13^. Saccharibacteria are strongly enriched in inflamed epithelial microbiomes which occur in gingivitis, periodontitis, bacterial vaginosis, and inflammatory bowel disease^10,14–19^; however, previous animal model studies have shown that Saccharibacteria can exert beneficial effects by reducing inflammation, particularly inflammation caused by their host bacteria^20^. This presents a complex question; are Saccharibacteria interacting with human cells directly to reduce inflammation, or are they reducing inflammation through effects applied to their host-bacteria? Understanding this question is essential to understanding the roles played by Saccharibacteria in disease initiation and progression. The interactions between Saccharibacteria and human oral epithelial cells remain completely uncharacterized, and studies on their Actinomycetia host-bacteria are also limited.

Saccharibacteria are characterized by their ultrasmall cell size (200–500 nm) and reduced genomes, which lack essential biosynthetic pathways, making them entirely dependent on other microorganisms for survival^4,21–24^. The first cultured Saccharibacterium, *Nanosynbacter lyticus* (*N. lyticus*) strain TM7x, grows on its cognate host *Schaalia odontolytica* (*S. odontolytica*) strain XH001, initially causing significant host death but eventually establishing stable symbiosis, even providing protection against phage infection in some cases^11,25,26^. Molecular investigation has identified two distinct type IV pili (T4P) systems in Saccharibacteria: one facilitating direct host-bacteria attachment and the other enabling twitching motility^27^. Despite their highly reduced genome, the retention of multiple T4P filaments suggests that T4P appendages play a crucial role in Saccharibacteria survival. While previous studies have shown that bacterial T4P can mediate interactions with human cells^28–32^, such a role has neither been demonstrated nor hypothesized for Saccharibacteria.

The human oral mucosal barrier, composed of multiple epithelial layers, is a key component of innate immunity and is constantly exposed to a wide range of stimuli, including polymicrobial communities, mechanical damage from mastication, and environmental antigens^33–36^. Numerous studies have established that oral epithelial cells express microbe-associated molecular pattern receptors (e.g., TLRs, NODs, PARs)^35,37–41^. These receptors initiate immune responses such as the release of proinflammatory cytokines to recruit monocytes, the secretion of antimicrobial peptides to regulate the commensal microbiome, and the activation of inflammasomes. Since Saccharibacteria and their host Actinobacteria are natural oral commensals, with 97% and 100% prevalence respectively^8,10,42,43^, they frequently interact with oral epithelial cells in healthy and diseased tissues. Early studies suggested that *Schaalia* and *Actinomyces* induce proinflammatory cytokines and antimicrobial defenses in gingival epithelial cells through TLR2 activation^44–48^. Additionally, cell wall components, particularly surface fimbriae and lipoproteins, have been shown to trigger inflammatory responses in neutrophils and macrophages^49–51^. *S. odontolytica* has also been identified as a major contributor to periodontitis and actinomycosis^44,45,52,53^ with emerging evidence suggesting roles in bacteremia and colorectal cancer progression^45,54,55^.

To address these knowledge gaps, our study aimed to directly characterize the interaction between Saccharibacteria—both alone and in association with its host bacterium—and oral gingival epithelial cells. These epithelial cells maintain homeostasis between the gingival microbiome and the mucosal barrier, influencing the development of gingivitis and periodontitis ^35,56,57^. Our findings reveal that *Saccharibacteria* alone do not induce significant proinflammatory cytokine production or gingival cell death. In contrast, we observed high levels of IL-8, MCP-1, and GRO-α secretion in response to the host Actinobacteria. Further analysis demonstrated that Saccharibacteria bind to gingival epithelial cells via a T4P-dependent mechanism, leading to clustering of TLR2 receptors and subsequent caveolin-mediated endocytosis. Internalization of *N. lyticus* results in low levels of TLR2 expression and surface availability, leading to dampening of cytokine responses during co-infection. Thus, Saccharibacteria are immune silent oral bacteria that directly associate with gingival cells to modulate inflammatory responses to other microorganisms, potentially contributing to microbial-immune system balance in the oral cavity.

## Results

### *N. lyticus* mitigates inflammatory cytokine response induced by its host bacteria

Bacterial interactions with oral epithelial cells were tested by exposing well-characterized human telomerase immortalized gingival keratinocytes (TIGK)^58–64^ cells to *N. lyticus type* strain TM7x alone, host bacteria XH001 alone, and a stable, well-established coculture of both, and measuring global cytokine response (81 cytokines) (Figure 1A)^11,65,66^. Prior to immune induction by TM7x, isolated episymbiont cells were extensively cleaned of host-bacteria materials via low-speed centrifugation and dialysis, preventing host antigen carryover (Figure S1A). Cleaned TM7x treatment displayed minimal IL-8 response compared to less cleaned treatments (undialyzed or high-speed centrifugation) (Figure S1B), while less cleaned XH001 supernatants (sup) induced a similar response as uncleaned episymbionts (Figure S1B). Cleaned TM7x size, viability, and quantity were characterized by live-dead fluorescence imaging, qPCR quantification, and NanoSight small particle analysis (Figures S1C-E), indicating that over 99% of cells remained viable after cleaning. Upon infection, XH001 cells strongly induced proinflammatory cytokines IL- 8 and GRO-α. TM7x and the host-episymbiont coculture induced a similar but reduced cytokine response for these mediators (Figure 1A). TM7x alone treatment induced exceptionally low IL-8 and GRO-α production, but higher TIMP-2 anti-inflammatory cytokine^67–69^ production (Figure 1A). To reinforce our results, we performed bacterial infection assays using two additional oral epithelial cell lines, normal oral keratinocytes-spontaneously immortalized (NOK-SI)^70,71^ and human gingival epithelial progenitor cells (HGEPp)^72–74^. Targeted quantification of IL-8 and GRO- α transcripts (RT-qPCR) and protein levels (ELISA) in TIGK, NOK-SI, and HGEPp showed similar results, strong inflammatory responses to XH001 treatment with significantly reduced responses from coculture and TM7x infections (Figures 1B-C, S2A, S2B). One additional inflammatory cytokine, MCP-1, was also highly induced in NOK-SI and HGEPp cells by XH001 and showed decreased pattern with TM7x and coculture (Figure S2A, S2C). These proinflammatory cytokines recruit inflammatory immune cells from the bloodstream, and IL-8 attracts neutrophils while GRO- α and MCP-1 attract monocytes and macrophages^75,76^. These results suggest TM7x does not induce strong innate immune response in oral epithelial cells, and can reduce cytokine responses induced by its host *S. odontolytica*. All infections resulted in almost 100% TIGK cell viability, so cytokine responses did not result from epithelial cell killing (Figure S2D).

**Figure 1.**
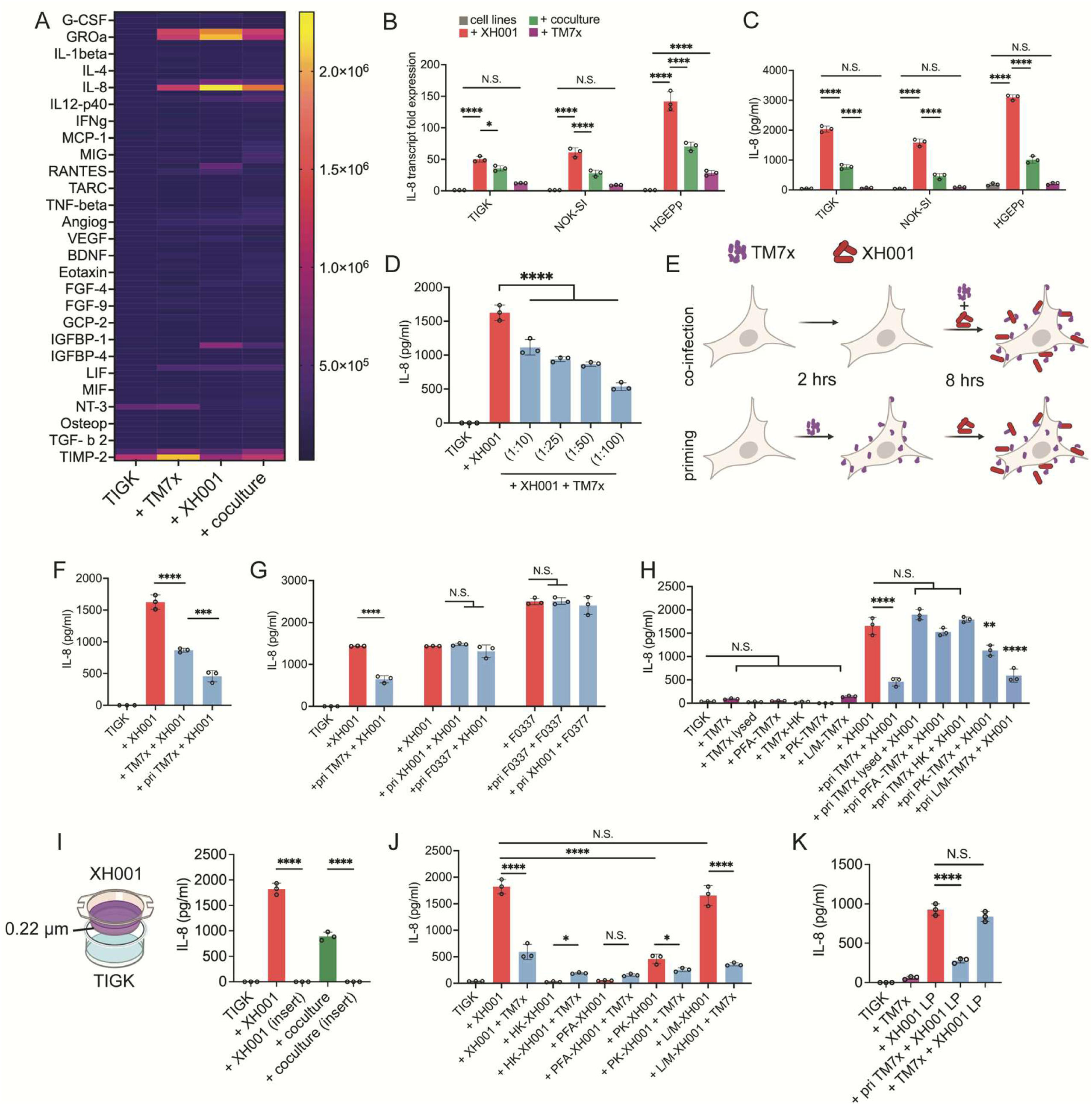
Cytokine response in oral epithelial cells. (A) Heatmap showing expression of 81 human cytokine proteins (RayBio array) for TIGK cells incubated with TM7x alone, XH001 alone or TM7x/XH001 coculture. The signal units are arbitrary grey value from the membrane array imaging. (B-C) Transcript fold change (B) and protein quantification via ELISA (C) of IL-8 in multiple epithelial cell lines infected with sham (grey), XH001 (red), TM7x (purple) and coculture (green) treatments. (D) TIGK cells (grey) treated with a fixed amount of XH001 (red) plus increasing concentration of TM7x at XH001:TM7x ratios of 1:10, 1:25, 1:50 and 1:100 (blue bars). (E-F) Graphical representation (E) and IL-8 expression (F) of TIGK cells that were either infected with XH001 and TM7x simultaneously or primed with TM7x before XH001 infection. (G) Results of repeating the priming experiment using XH001 or Actinomyces strain F0337. (H) IL-8 expression in TIGK cells infected with treated TM7x bacteria (purple) or XH001 (blue). +pri PFA-paraformaldehyde, +pri HK-heat killed, +pri PK-protease K, +pri L/M-lysozyme/mutanolysin. (I-K) IL-8 expression in TIGK cells grown with spatially isolated XH001 cells in an insert chamber (I), TIGK cells grown with neutralized XH001 treatments (J) and TIGK cells grown with isolated XH001 lipoproteins (K). Plots depict five independent experiments, all with three biological replicates. Means were compared via one-way ANOVA with Bonferroni correction for multiple comparisons with * P<0.05, ** P<0.01, *** P<0.001, **** P<0.0001, and NS = not significant.

To confirm our findings, isolated TM7x and XH001 cells were added concurrently to TIGK cells in increasing concentrations, indicating a dose-dependent decrease in IL-8, GRO-α, and MCP-1 with increasing TM7x presence (Figures 1D, S2E, S2F). To interrogate if TM7x reduces cytokine induction by directly engaging with TIGK cells or altering host-bacteria, we primed TIGK cells with TM7x for two hours prior to XH001 infection (Figure 1E). Pre-treatment with TM7x decreased XH001 induction of IL-8, GRO-α, and MCP-1 levels even further than the concurrent addition of TM7x and hosts (Figures 1F, S2G-H). The observed priming effect was TM7x specific since priming with XH001 or another *Actinomyces* sp. strain F0337 did not show the same IL-8 dampening (Figure 1G). Adding TM7x before host-bacteria or alongside host-bacteria had no effect on XH001’s viability (CFU/ml) when incubated with TIGK cells (Figure S2I). Therefore, the observed priming effect suggests that TM7x directly interacts with the TIGK cells to alter subsequent immune responses. To understand if this effect requires live Saccharibacteria, we added TM7x cells that had been neutralized using various methods (mechanical lysing, paraformaldehyde fixation, or heat treatment). All neutralized treatments resulted in abrogation of the cytokine response, suggesting live TM7x is crucial for the observed effect (Figure 1H). Protease-treatment of TM7x partially inhibited IL-8 induction; however, mutanolysin and lysozyme treatments did not, suggesting that TM7x’s interaction with TIGK cells is likely protein-based rather than polysaccharide-based (Figure 1H). Although TM7x induced TIMP-2 in our initial global screen (Figure 1A), when we used a targeted ELISA-based assays to quantify TIMP-2, this difference was significant only in TM7x treated group but not with a large effect size (Figure S2J). Previous studies have shown that various human cell cultures induce proinflammatory cytokines in response to cell wall components from *Actinomyces* and *Schaalia*^44,45,51^. To determine if XH001 triggers cytokine responses via direct surface interaction or diffusible molecules, we incubated XH001 in a transwell system that separates the TIGK cells from XH001 with a selectively permeable 0.22 µm membrane (Figure 1I). In this physically separated chamber, XH001 no longer induced IL-8, suggesting that XH001 immune activation is contact-dependent in TIGK cells (Figure 1I). Neutralizing XH001 via heat-killing, paraformaldehyde fixation, or protease K digestion reduced IL-8 induction, however lysozyme/mutanolysin treatment had no effect on IL-8 (Figure 1J), suggesting that the antigenic molecule is protein-based. A previous study^51^ suggested that cell surface lipoproteins on *A. viscosus* induced a cytokine response and XH001 also belongs to same genus Actinomyces, so we isolated the lipoproteins from XH001 and showed that they robustly induce IL-8 and that priming TIGK cells with TM7x decreases lipoprotein-dependent induction (Figure 1K). Simultaneous addition of TM7x and XH001 lipoproteins did not decrease cytokine response, possibly suggesting better competition and potency of isolated molecules.

### Epithelial transcriptome reveals activation of innate immune and vesicle trafficking pathways

To reveal oral epithelial genes and functions modulated by Saccharibacteria and their host-bacteria, we conducted global transcriptome analyses of TIGK cells infected with TM7x, XH001, and an established coculture. Our sequencing resulted in a total of 32,967,194 to 49,374,883 million reads per sample. 95.4-99.7% of transcripts were from TIGK cells, while the remaining transcripts were from the infecting bacteria. This aligned with expectation as bacteria produce fewer mRNA transcripts^77–79^. We recovered an average of approximately 2 million reads from TM7x in TM7x monoinfected cells, while an average of approximately 250,000 reads from XH001 monoinfected cells were recovered, suggesting TM7x may be actively transcribing within epithelial cells. Downstream analysis focused on epithelial transcripts because low bacterial reads prevented detection of differentially regulated bacterial genes. Principal component analysis (PCA) of gene expression across the four groups revealed that replicates within each group clustered closely in the space defined by the first three principal components, indicating high within-group similarity (Figure 2A). While the XH001-alone and TM7x-alone groups appeared closer to each other on the PC1 vs. PC2 plot, the three-dimensional PCA including PC3 showed clear separation among all four groups, highlighting distinct gene expression profiles. Notably, along PC3, XH001-alone and TM7x-alone were positioned far apart, whereas the coculture group occupied an intermediate position. This suggests that coculture induces a gene expression shift in XH001 group toward a TM7x-like profile. The top 200 most variable genes consistently showed that XH001 infection induced the largest difference compared to the TIGK cells alone treatment (Figure S3A, Table S1). We also observed many differentially regulated genes that were upregulated regardless of which bacterium was introduced. Species-specific and dual infection-specific genes comprise a small proportion of the upregulated genes detected (1,171 for XH001 vs. Control, 671 for TM7x vs. Control, and 322 for XH001+TM7x vs. Control) (Figures S3B-E). This suggests that there is a generalized bacterial response that is activated by bacteria across a wide phylogenetic range. Infection by XH001 induced more genes to be upregulated than infection by TM7x or by the host-episymbiont coculture, suggesting that XH001 induces a specific epithelial response that TM7x can suppress.

**Figure 2.**
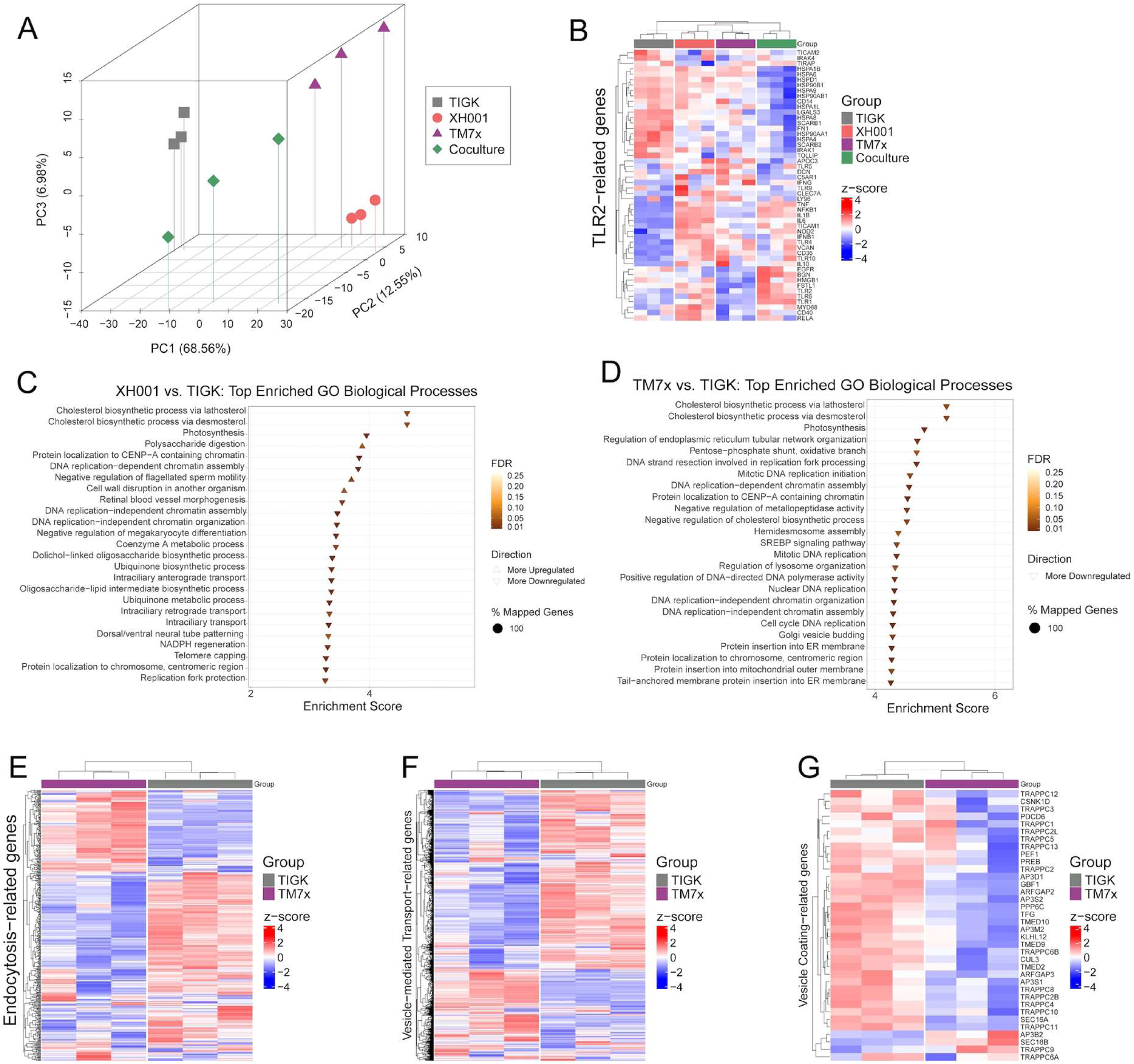
Global gene response of TIGK cells exposed to XH001 and TM7x. (A) Principal component analysis of PC1, 2 and 3 showed that all replicates within groups were clustered together while four groups had distinct clustering from each other. (B) Differentially expressed TLR2-related genes were plotted using heatmap with hierarchical clustering of both columns (samples) and rows (genes). Mirroring what observed in the PCA plot, uninfected cells cluster distinctly from all infection groups, monoinfection (XH001, TM7x, and coculture) groups cluster distinctly from the dual infection group. The dual infection group shows several genes being downregulated compared with the other treatment groups, including multiple heat shock proteins. (C-D) GO analysis for biological function show upregulated and downregulated biological process in TIGK compared with XH001 infected TIGK cell (C) and between TM7x infected TIGK (E-G) RNASeq data were normalized via logarithmic transformation and plotted to visualize pathways for endocytosis (E), vesicle-mediated transport (F), and vesicle coating (G). Hierarchical clustering on columns (samples) and rows (genes). Mirroring what is found in the PCA plot, the TIGK cells only distinctly cluster from TM7x infected groups indicates similar trends. Infection by TM7x had unique effects on endocytosis, vesicle-mediated transport, and vesicle coating processes.

A closer look at the expression changes induced by XH001 revealed upregulation of many innate immunity genes (Figure S3C, Table S1). Relative to TM7x infection alone, we observed that infection by XH001 and the coculture induced upregulation of TLR2 and TLR2-related pathways (such as TLR6, TLR1, MyD88, IL-6, IL-8, IL1-β, and NF-κB) (Figure 2B). This supports previously published results showing TLR2-dependent immune activation by Actinomyces^44–48^. However, infection by the episymbiont-host coculture induced a significantly weaker immune response relative to host bacteria alone. To understand the regulatory pathways modulated by TM7x, we identified enriched functional gene clusters using STRING^80^ and graphed the most impacted pathways (GO term-derived Biological Processes)^81^. Treatment with XH001 alone impacted chromatin remodeling, DNA replication, and cell proliferation genes (Figure 2C), suggesting an alteration in TIGK replication and cell division. In contrast, TM7x modulated multiple pathways for vesicle/membrane trafficking, endoplasmic reticulum processing, and endocytosis-related genes (Figure 2D). Examining functional genes within these vesicle/membrane-related pathways revealed that TM7x broadly downregulated expression relative to TIGK cells alone (Figures 2E-G, S3F-H). Examples included SCARB2, which is involved in the biogenesis and maintenance of endosomes; GRN, which regulates endocytosis; BECN1, TMEM106, and TREX1, which regulate early and late endosomes; and ARL6IP1, which facilitates ER trafficking. These results suggest XH001 induces cytokine responses through the TLR2 pathway while TM7x potentially modulates endocytosis and vesicle trafficking pathways.

### Epithelial TLR2 pathway mediated response to *S. odontolytica* and *N. lyticus*

To validate our transcriptomic findings, we employed α-TLR2 antibody to block TLR2- mediated immune responses in TIGK cells. α-TLR2 antibody treatment completely neutralized TIGK responses to both XH001 and TM7x-XH001 coculture infection, suggesting that TLR2 is an essential receptor for the epithelial cytokine response induced by XH001 (Figure 3A). For additional support, HEK293-Null and HEK-TLR2 expressing cell lines were infected with XH001, TM7x, and the coculture, with or without α-TLR2 antibody. In HEK293-Null cells (which do not express TLR2) IL-8 production was not induced by any of the infections (Figure 3B). HEK293- TLR2 cells produced elevated IL-8 when infected by XH001 or the coculture, and that response was completely inhibited by addition of α-TLR2 antibody (Figure 3A), showing that expression of TLR2 was sufficient to confer XH001 induced IL-8 production. IL-8 induction by TM7x in HEK293- TLR2 cells was low but significant, suggesting that TM7x also interacts with TLR2 to induce low-level immunoactivation. Consistent with the latter idea, HEK293-TLR2 cells primed with TM7x still inhibited XH001 responses (Figure 3B), suggesting that TM7x-dependent inhibition is acting through TLR2-dependent processes. To confirm this, we induced IL-8 expression in TIGK cells using TLR2-targeting agonists Pam2CSK4 (specific to TLR2/TLR6 complexes) and Pam3CSK4 (specific to TLR2/TLR1 complexes)^82–84^. TIGK cells were activated by both agonists, but TM7x priming only inhibited activation by Pam3CSK4 (Figure 3C), suggesting that TM7x not only interacts with TLR2, but prefers the TLR2/TLR1 heterodimer.

**Figure 3.**
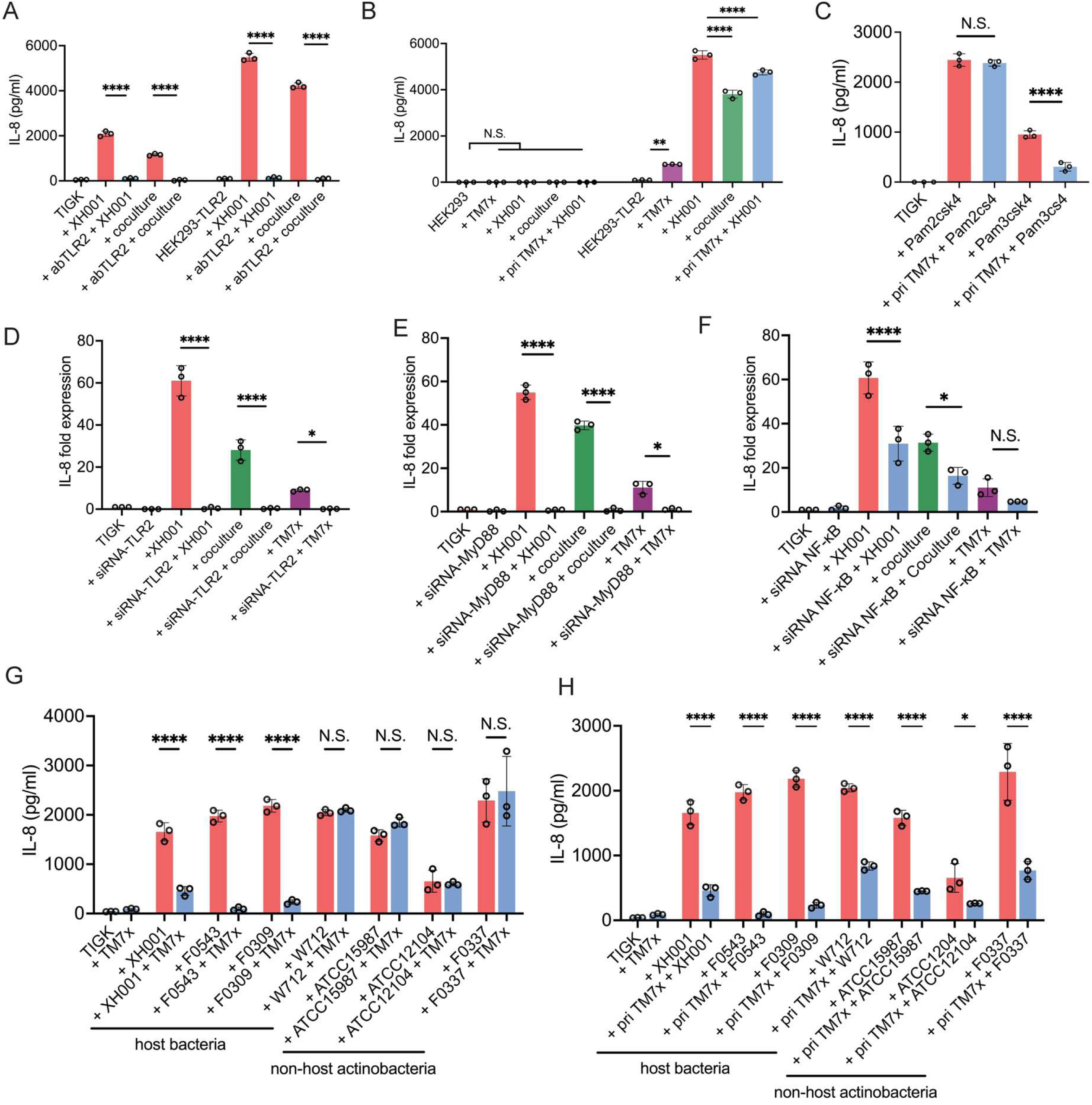
XH001 induces cytokines via TLR2-dependent pathways which are inhibited by TM7x. (A) TIGK and TLR2 expressing HEK293 cells (HEK293-TLR2) were treated with α-TLR2 antibody before treatment with XH001 and TM7x/XH001 cocultures. (B) XH001 (red), coculture (green), TM7x (purple), and TM7x priming followed by XH001 infection (blue) of HEK293-TLR2 and HEK293-Null cells. HEK293-Null cells had a minimal response while TLR2 expressing cells had a TIGK-like response. (C) TIGK cells primed with TM7x (blue) were treated with TLR2/TLR6 agonist pam2CSK4 and TLR2/TLR1 agonist pam3CSK4 (red). (D-F) TLR2, MyD88, and NF-κb were knocked down in TIGK cells using siRNA. For the quantification gene silencing, see Figure S4. si-RNA treated cells were infected with XH001 (red), coculture (green) and TM7x (purple) then IL8 cytokine expression was quantified. (G-H) TIGK cell IL-8 response to various Actinobacteria strains with (blue) and without (red) TM7x were quantified. TM7x was either added at the same time as the host bacteria (G) or added before the host bacteria (H). Three biological replicates were completed for every experiment. Means were compared using one-way ANOVA with Bonferroni correction for multiple comparisons with * P<0.05, ** P<0.01, *** P<0.001, **** P<0.0001, and NS = not significant.

To elucidate the mechanism of immunoactivation, TLR2 and two downstream TLR2- pathway genes, MyD88 and NF-κB, were knocked down via small interference RNA treatment (siRNA) in TIGK cells^85–88^. We designed siRNAs against TLR2, MyD88, and NF-κB that inhibited target gene expression by >90% and used scrambled RNA as a negative control (Figures S4A-E). Knockdown of TLR2 decreased induction of IL-8 and GRO-α in all bacterial infection treatments, and knockdown of MyD88 and NF-κB decreased IL-8 in the same groups (Figures 3D-F, S4F). This suggests that TLR2 and its downstream induction pathway are crucial TIGK cells to mount a cytokine response to XH001, and also for TM7x to inhibit the observed immunoactivation. To broaden these findings, we tested both episymbiont-host and non-host Actinobacteria related to XH001^20,89^ (Key Resources Table). All tested Actinobacteria induced high levels of IL-8, GRO-α, and MCP-1. Concurrent addition of TM7x alongside compatible host bacterial species inhibited pathway induction, but did not inhibit activation by non-host Actinobacteria (Figures 3G, S4G, S4I). Preemptively priming TIGK cells with TM7x decreased pathway induction by all tested Actinobacteria to similar degree (Figures 3H, S4H, S4J). Similar to XH001, cytokine induction by these Actinobacteria was TLR2-dependent, illustrated by significantly reduced cytokine production when TLR2 was knocked down via siRNA (Figures S4K-L). This suggests that tested Actinobacteria can induce epithelial immunity via the TLR2 pathway, and TM7x can effectively inhibit this induction given sufficient time to interact with the epithelial cells.

### Type IV pili and TLR2 mediated *N. lyticus* binding to oral epithelial cells

TM7x priming and TLR2 inhibition experiments suggested that TM7x directly interacts with TIGK cells, potentially via contact-dependent binding. To visualize potential binding, TM7x with and without MitoTracker fluorescent labeling was added to TIGK cells. MitoTracker dye^90^ stains bacterial membranes based on electrostatic potential^91,92^ and does not impact TM7x viability or infectivity (Figures S5A-B). With or without the MitoTracker dye, TM7x cells showed robust binding to TIGK cell surfaces relative to the untreated control (Figures 4A-B, S5C-H). Focal stacking in the z-axis revealed TM7x was detectable diffusely across TIGK cells (Figures S5I-J). Flow cytometry quantification of MitoTracker labeled TM7x validated the observed TIGK cell binding. Remarkably, ∼100% of screened TIGK cells were associated with TM7x, while TIGK cells inoculated with similarly sized inert resin beads (∼500 nm) showed only 2% bead-binding, suggesting that the observed binding is specific to the TM7x and not a result of its small cell size (Figures 4C-D). Fluorescence-labeled TM7x cells that had been neutralized by paraformaldehyde fixing, ethanol treatment, or mechanical lysis did not bind TIGK cells (Figure 4E), suggesting that TM7x binding required intact, living Saccharibacteria. Longitudinal quantification revealed that TM7x binding is rapid and efficient, becoming detectable ∼15 minutes post-infection, and reaching saturation ∼ 2 hours post-infection (Figures S5K-L). Consistent with our cytokine analyses, MitoTracker-TM7x showed robust binding to multiple oral epithelial cell lines (NOK-SI, HGEPp, OKF-6^93^) but not the HEK293-Null cells that lack TLR2 (Figures 4F, S6A-D). We also tested the binding of XH001, coculture, and additional Saccharibacteria strains to TIGK cells using the same method. Coculture had ∼76% binding, while XH001 had the lowest ∼46% binding, suggesting that TM7x interacts with oral epithelial cells more robustly than its host bacteria XH001 (Figure S6E). Saccharibacteria strains BB002, BB004, and BB008 (all originally isolated from periodontal patients samples^20,89^) exhibited strong binding (∼82-99%), however *Southlakia epibionticum*^94^ (originally isolated from saliva samples) showed significantly lower TIGK binding (∼20%). Multiple, but not all, Saccharibacteria strains bind oral epithelial cells and it appears to be most common amongst strains that reside in periodontal pockets (Figure S6E).

**Figure 4.**
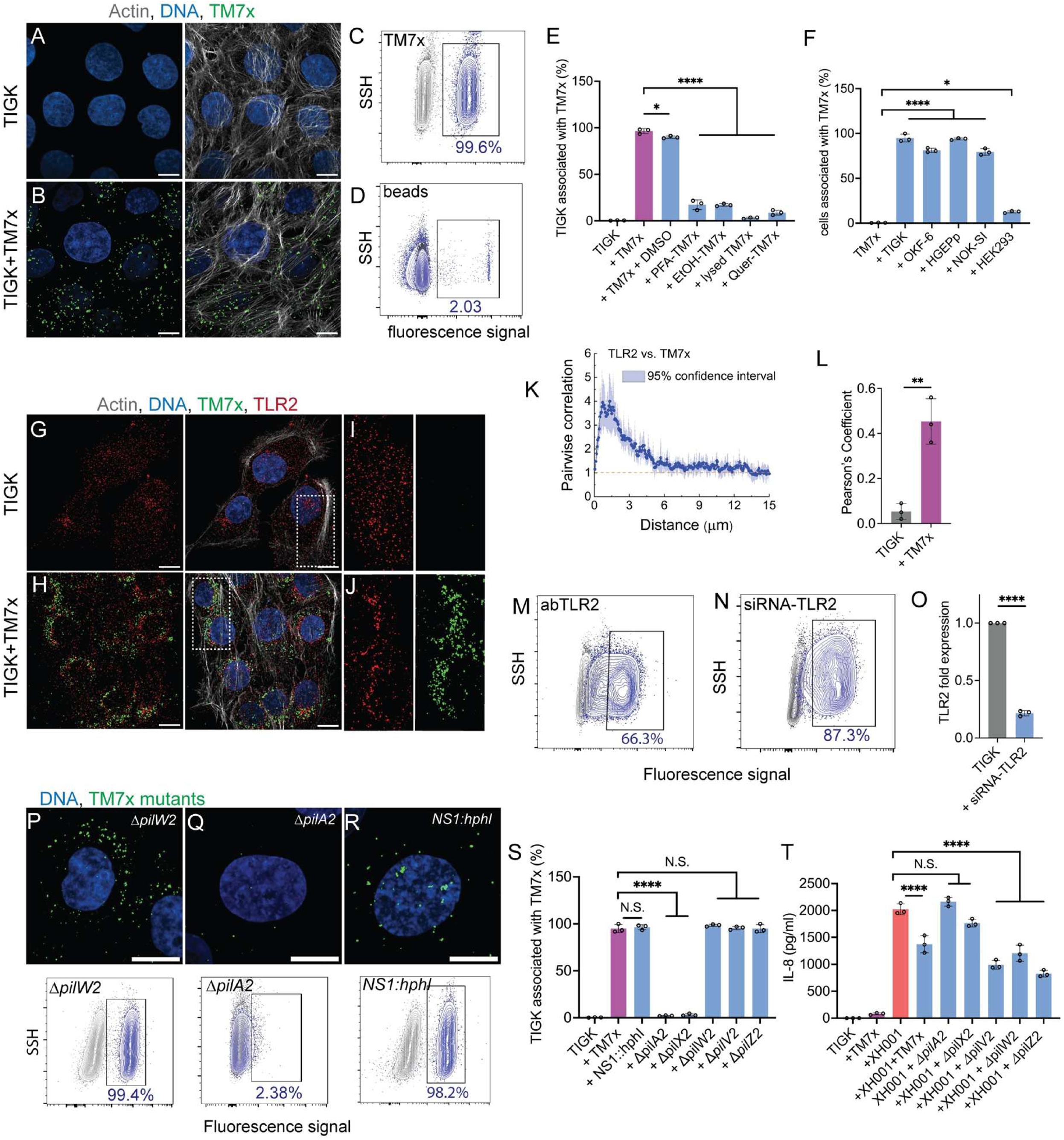
TM7x attachment to TIGK cells through type IV pili and TLR2. (A-B) MitoTracker stained TM7x cells (green) were visualized by confocal microscope after infecting TIGK cells. Actin cytoskeletons and nuclei were stained with phalloidin (grey) and DAPI (blue) respectively. (C-D) TIGK cells bound to MitoTracker-TM7x (C) or ∼500 nm size beads (D) were quantified using flow cytometry. TIGK cells alone (grey) were similar to TIGK cells exposed to small beads (blue), but distinct from cells bound by TM7x (blue). (E) Treating TM7x with paraformaldehyde (PFA), ethanol (EtOH), physical lyses, or quercetin reduced epithelial cell binding as quantified by flow cytometry. (F) Flow cytometry analysis demonstrated MitoTracker-TM7x binding various oral epithelial cell lines. (G-J) Co-staining infected TIGK cells for TM7x-MitoTracker (green) and TLR2 immunofluorescence (red) shows co-localization, actin (grey) and DNA (blue) stained to visualize intracellular structure. (I) and (J) are zoomed-in insets from (G) and (H) respectively. (K-L) daime and Pearson’s coefficient analysis of TIGK cells co-immunostained for TM7x and early endosome marker EEA1. (K) daime analysis for images in (G) and (H) demonstrated positive correlation values between TLR2 and TM7x all the way through a dipole distance of ∼10 µm, which indicated the close spatial proximity relationship across the epithelial cells. (L) Pearson’s coefficient analysis for the same images show strong colocalization of TM7x and TLR2 receptor. (M-N) Flow cytometry of MitoTracker-TM7x binding TIGK cells in the presence of α-TLR2 antibody or TLR2 si-RNA knockdown cells. (O) Quantification of TLR2 transcripts in TIGK cells treated with α-TLR2 siRNA. (P-S) Example fluorescence images and flow cytometry analysis of TIGK cells infected with MitoTracker-TM7x T4P mutants. Complete data for T4P mutant strains found in Figure S6. NS1::HphI indicates a neutral mutant that has the hygromycin B resistance marker inserted into a validated neutral site to provide a negative control. (T) T4P mutants defective for T4P-2 function (Δ*pilA2* and Δ*pilX2*) could not effectively suppress XH001-dependent induction of IL-8. Three biological replicates were completed for every experiment. Means were compared using one-way ANOVA with Bonferroni correction for multiple comparisons with * P<0.05, ** P<0.01, *** P<0.001, **** P<0.0001, and NS = not significant. All scale bars are 10 µm.

Previous studies have found some bacteria that can directly bind surface TLR2^87,95,96^. To determine if Saccharibacteria directly bound TLR2, we co-stained TM7x-MitoTracker and TLR2 immunostaining in TIGK cells and calculated colocalization via Pearson’s coefficient^97^ and digital image analysis in microbial ecology (daime) proximity analysis^98^. This revealed that TM7x closely colocalized with TLR2 and induced significant TLR2 clustering near the nucleus, while uninfected TIGK cells showed no TLR2 clustering and no antibody cross-reactivity (Figures 4G-L, S6F-G). Treatment of TIGK cells with TLR2 neutralizing monoclonal antibody reduced TM7x binding in flow cytometry experiments by ∼33% and treatment with TLR2-siRNA reduced binding by ∼12.7%. This partial inhibition could be due to the fact that both antibody and siRNA do not neutralize 100% of the receptors^85,99,100^. These data suggest TLR2 is not only important for modulating TM7x-dependent cytokine inhibition, it also a putative binding substrate (Figures 4M-O).

Type 4 pili (T4P) are broadly conserved surface appendages that are known to adhere to epithelial tissues in other model bacteria^30,32,101,102^, are capable of binding TLR2^103,104^, and are enriched in bacterial episymbiont genomes^13,27^. Additionally, comparing T4P gene expression between TM7x-XH001 cocultures and TM7x alone in the presence of TIGK cells revealed that most T4P genes were expressed at similar or even higher levels in the TM7x-alone condition, suggesting that T4P may play an important role in TM7x-TIGK interaction (Figure S6H). To determine if TM7x was using T4P to bind TIGK cells, we added the non-specific T4P chemical inhibitor quercetin to the assay^13^. Quercetin induced a ∼97% decrease in TM7x binding (Figure 4E). Our previous study showed that clade G1 Saccharibacteria, including TM7x, produce two functionally distinct filaments, T4P-1 and T4P-2^27^. T4P-1 provides twitching motility and is essential for cell survival. T4P-2 facilitates host bacterial binding at a distance and mutants have been constructed for most of its subunits. To determine if T4P-2 impacts TIGK binding, cells were infected with either T4P-2 producing strains (Δ*pilW2,* Δ*pilV2,* and Δ*pilZ2*) or T4P-2 deficient strains (Δ*pilA2* and Δ*pilX2*). Loss of the PilA2 major pilin (∼2%) and PilX2 minor pilin (∼3% bound) drastically reduced TIGK binding, and all T4P-2 producing mutants^27^ showed no change in binding (Figures 4P-S, S6I-L). This was confirmed by both confocal imaging and flow cytometry quantification. Furthermore, induction of IL-8 expression by XH001 was significantly inhibited by co-infection with all TM7x mutants except Δ*pilA2* and Δ*pilX2* (Figure 4T), confirming that T4P-2, and presumably direct cell-to-cell binding, is essential for TM7x to successfully modulate immunoactivation of oral epithelial cells.

### Caveolin-dependent internalization of Saccharibacteria reduced TIGK response

Analysis of individual z-stacks within confocal images (Figure S5J) revealed that the majority of TM7x cells appeared inside TIGK cells, suggesting that TM7x may invade, or get internalized by, TIGK cells. This is also supported by our transcriptomic data where many of the endocytosis and vesicle trafficking genes were differentially regulated. To investigate, TM7x- reactive polyclonal antibodies were generated (see methods) and used to differentially stain extracellular and intracellular TM7x cells (Figure 5A) via cell permeabilization and sequential immunostaining. Confocal imaging indicated distinct populations of extracellular and intracellular TM7x. This suggests that after binding to surface TLR2, TM7x cells can become internalized within TIGK cells (Figures 5A, S7A). Quantification of images revealed that after 8 h post-infection, ∼67% of detected TM7x were intracellular and ∼33% were extracellular (Figure S7B). Transmission electron microscopy (TEM) was used to visualize infected TIGK cells. TM7x was clearly visible on both cell surfaces and intracellularly, encased in endosomal vesicle-like structures throughout the cell body that were absent in uninfected TIGK cells (Figures 5B-E). These endosomal vacuole-like structures with enclosed TM7x had a clear electron-lucent zone between the membrane and TM7x cells, suggesting TM7x may exude a thick polysaccharide layer or glycocalyx as seen in many pathogenic bacteria^105–108^ (Figures 5F-H). Additionally, some vesicles contained both intact and compromised TM7x cells with releasing cytoplasmic content (Figures 5I-K), indicating that these structures are possibly early and late endosome compartments where foreign particles are neutralized over time via lysosomal enzymes^109,110^.

**Figure 5.**
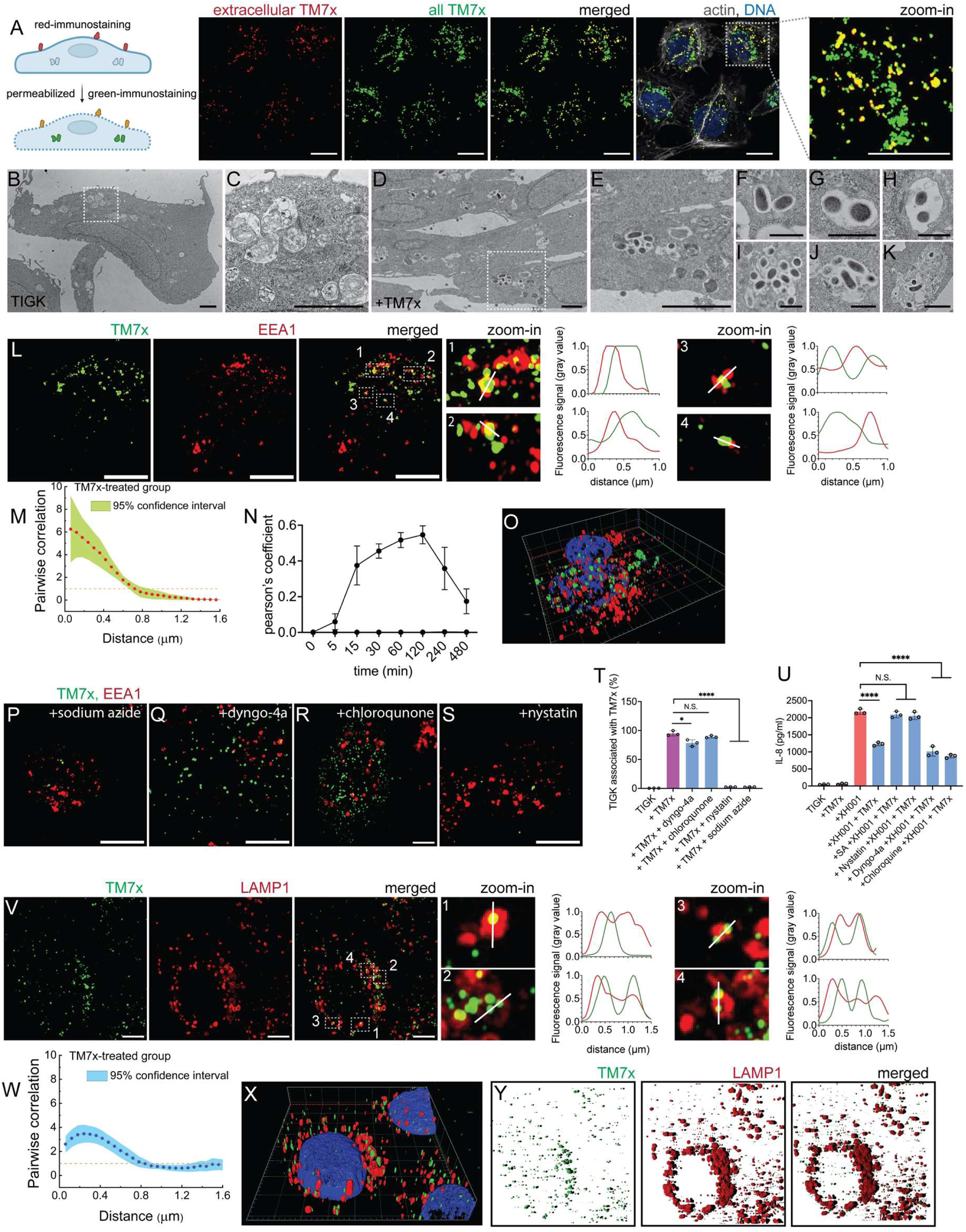
Endocytosis of TM7x by TIGK cells. (A) TM7x infected TIGK cells were differentially stained by applying one secondary antibodies before cell permeabilization (red) and another after cell permeabilization (green), resulting in extracellular TM7x appearing yellow and intracellular TM7x appearing green. DNA was stained with DAPI and F-actin was stained with phalloidin. All scale bars are 10 µm. (B-K) Transmission electron microscope images of uninfected TIGK cells (B-C) and TM7x infected cells (D-K). Panels (C) and (E) are zoomed-in images from panels (B) and (D). Scale bars are 2 µm (B-E) or 0.5 µm (F-K). (L) Co-immunostaining of EEA1 (red) and TM7x (green) in infected TIGK cells. L1-L4 are zoomed-in area shown with white boxes (dashed lines), focusing on close proximity staining of EEA1 and TM7x. Inset graphs show the fluorescence intensity of the EEA1 and TM7x staining cross distance that is illustrated by the while lines. (M-N) daime and Pearson’s coefficient analysis of EEA1 and TM7x co-localization. (M) daime analysis for demonstrated strong pairwise correlation value at short distances (< 0.8 um) indicated that EEA1 is consistently adjacent to, or overlapping with, TM7x cells. (N) Pearson’s coefficient analysis of a time series indicates that endosome co-localization begins ∼15 minutes post-infection and increases until ∼2 hours post-infection. (O) 3D reconstruction of panel L, illustrating how TM7x cells colocalize with EEA1. (P-S) Co-immunostaining of EEA1 (red) and TM7x (green) were done on TIGL cells that were infected with TM7x in presence of various indicated endocytosis pathway inhibitors. (T-U) Quantification of MitoTracker-TM7x binding using flow cytometry and IL-8 response using ELISA quantification in the presence of endocytosis inhibitors. (V) Co-immunostaining of late endosome marker LAMP-1 (red) and TM7x (green) in TIGK cells. V1-V4 are zoomed-in boxes focusing on close proximity staining of LAMP-1 and TM7x. Inset graphs show fluorescence intensity of LAMP-1 and TM7x cross distance that is illustrated by the while lines. (W-Y) daime and 3D reconstruction analysis as shown in panels M- O of LAMP-1 and TM7x staining. Panel W show the daime analysis for images in V panel and additional replicates demonstrated the same phenomenon like EEA1 analysis with high positive pairwise correlation value at shorter distances (< 0.8 um) indicated that LAMP-1 are closely adjacent to TM7x cells, even overlapping. Panel Y shows each fluorescence channel separately in 3D reconstruction to highlight that most of the green TM7x stains are within the red LAMP-1 stain. Three biological replicates were completed for every experiment. Means were compared using one-way ANOVA with Bonferroni correction for multiple comparisons with * P<0.05, ** P<0.01, *** P<0.001, **** P<0.0001, and NS = not significant. All scale bars in panels A, L, P, Q, R, S, V are 10 µm.

To detect active endocytosis of TM7x by TIGK cells, TM7x cells were co-immunostained with early endosome antigen 1 (EEA1), a common marker protein for early endocytosis^111,112^. TM7x naturally colocalized with, or near, EEA1 staining (Figure 5L). EEA1 proteins are not a membrane maker, so they typically appear next to endocytosed particles rather than surrounding them^111,113^, as we observed with TM7x cells (Figures 5L1-4). Daime analysis predicted a strong association of EEA1 and TM7x within 0-0.7 µm distance (Figure 5M), while temporal Pearson’s coefficient tracking revealed that internalization starts 15 minutes post-infection and peaks 1-2 hours post-infection (Figure 5N). 3D rendering of confocal images confirmed the colocalization and adjacency of the immunostained EEA1 and TM7x (Figures 5O, S7C-D).

Endocytosis in epithelial cells is dependent on clathrin-dependent, dynamin-dependent, caveolin-dependent, and/or micropinocytosis pathways^114^. To inhibit these pathways, we first treated TIGK cells with the global metabolic inhibitor sodium azide to block all endocytosis events^115^. Sodium azide-treated cells failed to bind or internalize TM7x in both fluorescence imaging and flow cytometry (∼2%) experiments (Figures 5P, 5T). To interrogate specific pathways, we tested three additional inhibitors – chloroquine^116^, Dyano-4a^117^, and Nystatin^118^ – that inhibit clathrin, dynamin, and caveolin, respectively. Of these inhibitors, only nystatin treatment reduced caveolae formation, prevented colocalization and impaired TM7x binding with EAA1 (∼3% bound) (Figures 5Q-T). Chloroquine (∼92% bound) and Dyano-4a (∼85% bound) treatments had no significant effect. Additionally, only nystatin and sodium azide treatments ameliorated TM7x-dependent cytokine inhibition (Figure 5U). These findings indicate that TIGK cells rely on caveolin-dependent endocytosis to internalize TM7x.

Antibodies specific for a late endosome marker protein, lysosomal associated membrane <u>p</u>rotein-1 (LAMP-1), were co-stained alongside TM7x to track processing of internalized cells. LAMP-1 is located on membranes from late endosomes and lysosomes and would be expected to surround encapsulated materials^119–121^. TM7x and LAMP-1 demonstrated colocalization 4-8 hours post-infection while many internalized TM7x were surrounded by the LAMP-1 containing membranes (Figure 5V). We also observed TM7x staining on these membranes and even outside the periphery of the vesicles (Figures 5V1-4). Daime analysis predicted a strong association of LAMP-1 and TM7x within 0-0.8 µm distance (Figure 5W). Unlike EEA1 co-staining, 3D reconstruction with LAMP-1 staining revealed that most TM7x were engulfed by the LAMP-1 containing membranes (Figures 5X-Y, S7E-F), presumably inside the endosomes or lysosomes. To investigate whether endocytosed TM7x escaped into the cytosol and/or proceeded into autophagosomes for degradation, we co-stained infected TIGK cells with TM7x and the autophagosome marker protein 1a/1B light chain 3 (LC3)^122,123^. We did not observe colocalization of LC3 and TM7x, demonstrating that endocytosed TM7x does not transition into autophagosomes (Figure S7G).

### Endocytosed TM7x survive within oral epithelial cells and retain the ability to reinfect host bacteria

We investigated whether intracellular TM7x remained viable during endocytosis using a TIGK cell antibiotic protection and reinfection assay. TIGK cells were infected with MitoTracker- TM7x and then washed and treated with a combination of antibiotics (gentamicin and neomycin) to eliminate any extracellular TM7x (Figure 6A). Experimental validation of these antibiotics illustrated their ability to completely kill TM7x in the absence of TIGK cells (Figure S7H). Endocytosed TM7x were incubated for 8, 24, and 48 hours to track survival over time. After antibiotic treatment, TIGK cells were lysed, and lysates were added to XH001 cultures to assess the liberated TM7x’s ability to infect and grow (Figures 6B-F). At all-time points tested, viable TM7x cells were recovered and were capable of infecting XH001. However, extended intracellular incubation significantly reduced the viable cells recovered. Quantification of TM7x DNA using qPCR revealed a slow but continuous decrease in TM7x presence (Figure 6G). Within the context of human oral surfaces, epithelial cells undergo numerous physical and chemical challenges within a 48 hour period^35,124^. Epithelial cells can naturally or physically lyse by desquamation, chewing, speaking and oral hygiene practices, releasing cytoplasmic content back into the microbiome^125,126^. Even if TM7x cannot kill and lyse the TIGK cells, they could reasonably persist within cells and escape when natural or damage induced cell deaths occurs in the human oral cavity^127,128^. To explore this possibility, we wanted to find an upper limit on TM7x’s intracellular survival by extending post-infection incubation times. Infected TIGK cells were incubated for 4 days before being passaging into new fresh medium. This was done for three consecutive passages, equating to 8-10 generations in TIGK cells. At each passage, we observed decreasing amounts of TM7x according to fluorescence imaging, while differential staining of TM7x showed presence of only intracellular TM7x in passage 1 and passage 2 (Figures 6H-K), host re-infection assays, and qPCR analysis (Figures 6L-M). By passage three, TM7x was essentially undetectable. We concluded TM7x can survive multiple days (∼8 days) inside TIGK cells “waiting” for a natural/induced cell lysis event to enable reinfection of host bacteria, but eventually they are killed by late endosome and lysosome degradation.

**Figure 6.**
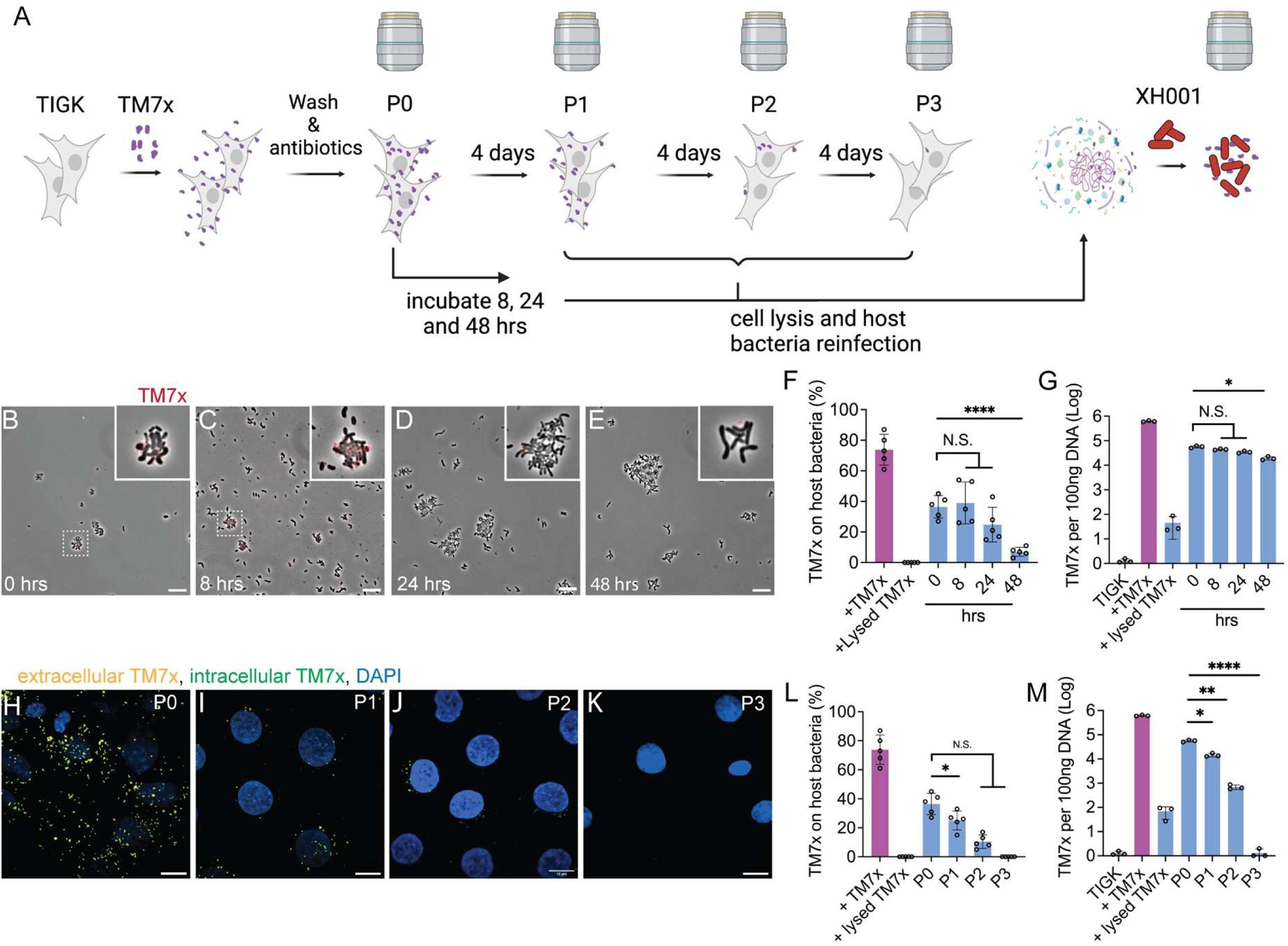
Persistence of TM7x within TIGK cells. (A) Experimental procedure for testing TM7x’s viability and after endocytosis within TIGK cells. After TM7x infection and endocytosis, TIGK cells were washed and treated with antibiotics to kill extracellular TM7x. Cell cultures were then incubated 8 hours, 24 hours, or 48 hours before passaging (three passages with a four**-**day interval). At multiple time points, TIGK cells were imaged for TM7x, qPCR quantified for TM7x, or lysed and added to XH001 to test TM7x viability/infectivity. P0-P3 indicate passages 1-3. (B-E) Phase contrast images of MitoTracker-TM7x recovered and infected XH001 from lysed TIGK cells after 8, 24, and 48 hours of endocytosis. MitoTracker-TM7x can be clearly at time 0 but decrease at later time points. (F-G) Quantification of recovered TM7x cells via XH001 re-infection (F) and qPCR indicate prolonged persistence intracellularly (G). To quantify infection rate in panel F, 60-100 XH001 cells per image for 5 images were quantified for attachment of TM7x on each cell. (H-K) Extra and intracellular localization of TM7x in TIGK cells visualized similar to in Figure 5A. After each passage, TIGK cells continued to grow while the number of TM7x cells decreased. By passage 3, we did not detect any TM7x staining. (L-M) Quantification of recovered and infected TM7x cells on XH001 (M) and qPCR quantified TM7x gDNA in infected and lysed TIGK cells (M) after multiple passages. Means were compared using one-way ANOVA with Bonferroni correction for multiple comparisons with * P<0.05, ** P<0.01, *** P<0.001, **** P<0.0001, and NS = not significant. All scale bars are 10 μm.

## Discussion

Decades of research have linked polymicrobial communities from the oral microbiome to enamel loss, inflammatory disease progression, and systemic infections^129–131^. However, microbial "dark matter" organisms such as the Saccharibacteria have always been a gap in our knowledge due to the technical difficulties associated with cultivating and studying them^132–134^. We investigated how Saccharibacteria interact with the first line of innate immune defense to the oral microbiome, epithelial cells, revealing that these ultrasmall bacteria can actively modulate immune cell responses in a contact dependent manner. Oral Actinobacteria, such as TM7x’s host species, trigger the TLR2-MyD88-NF-kb signaling pathway with their cell surface lipoproteins, inducing strong immune responses in epithelial cells, including proinflammatory cytokine production (IL-8, GRO-α, and MCP-1)^51,88,135^. Incredibly, episymbiont Saccharibacteria can inhibit host-induced immunoactivation by directly interacting with surface TLR2 present on epithelial cells using type IV pili (T4P) appendages, resulting in endocytosis of Saccharibacterium and potentially TLR2. While endocytosed TLR2 can trigger additional immune responses, it can also inactivated TLR2^136^—a mechanism that parallels the TM7x-induced dampening of cytokine responses we observed. Upon internalization, Saccharibacteria are ultimately directed to lysosomal degradation and killing, however prolonged intracellular survival presents opportunities for these bacteria to escape via epithelial cell death. Thus, the Saccharibacteria lifecycle in the oral cavity likely alternates between colonizing host bacteria, which enables reproduction, and residing within epithelial cells, which provide a natural reservoir to protect against transient stressors. This alternating episymbiont – reservoir lifestyle could explain the extremely high prevalence^8,137^ of Saccharibacteria amongst the human population despite being obligate epibionts with limited host-bacteria ranges^66,89^ and biosynthetic capacity^10,138^. Their ability to “peacefully” interact with oral epithelial cells and dampen host-bacteria immunoactivation suggests a stabilizing role in regulating microbiome-immune interactions.

The gingival crevice occupied by these microbes is a sensitive site where epithelial lining becomes progressively thinner at the oral sulcular and gingival epithelium. These epithelia are constantly stimulated by oral bacteria, including pathobionts, such as *Actinomyces* and *Schaalia*^44,45,130,139^. We observed these two bacteria induced strong proinflammatory cytokine responses, presumably intended to recruit immune cells to the gingival crevice, and episymbiotic Saccharibacteria modulated that inflammatory response, whether added before Actinobacteria or simultaneously. We have previously shown that Saccharibacteria modulate their host bacterial cell surface molecules, making them less immunogenic^20^; however, this study clearly illustrated that Saccharibacteria can directly interact with epithelial cells to suppress cytokine response. We believe this indicates that Saccharibacteria actively interact with their hosts, Actinobacteria and human epithelial cells, in the oral cavity. Our data indicate immunomodulation can be species specific for cognate episymbiont-host pairs and that the diverse members of Saccharibacteria and Actinobacteria could play complex roles in innate immunity. For example, we observed TM7x inhibited immune activation by most host-species, but one particular host, *Actinomyces meyeri* strain W712, was not affected. Furthermore, all tested Saccharibacteria showed some TIGK cell association, but *S. episymbioticum* showed much less binding than the other strains. These variations could explain why some Saccharibacteria and their host-bacteria species are abundant in healthy tissues while others are abundant within diseased tissues and sites of inflammation^6,20,43–45,129,130,140^. Future characterization of additional Saccharibacteria species will likely reveal the range of immunological roles played by different Saccharibacteria species, within different human body sites.

Numerous bacteria can directly bind to epithelial cells and become endocytosed^141–143^. This is especially true for upper respiratory tract commensals and pathogens^144–147^. Some of these interactions are facilitated by direct binding between type I pili and TLR2 receptors^103,148,149^. Once endocytosed, some commensals are killed by lysosome fusion, while others have molecular mechanisms to neutralize lysosome or escape endosomes and reach the cytosol^109,150,151^. Amongst the spectrum of all endocytosed bacteria, Saccharibacteria are unique due to their small size (200-300 nm) and tendency to be endocytosed in aggregates. This endocytosis also aggregates cell surface TLR2 receptors, removing them from the cell surface and concentrating them within intracellular vesicles. This exciting mechanism of immunomodulation inhibits proinflammatory responses against host and non-host Actinobacteria given sufficient pre-exposure, suggesting that Saccharibacteria could potentially dampen immune response to any bacteria. Saccharibacteria occasionally detach from host-bacteria and assume a motile free- floating state^66^, allowing horizontal infection of new hosts. Our findings suggest that this free-floating lifestage is also essential for transient association with epithelia that act as an environmental reservoir. One potential application of this mechanism could be the therapeutic addition of immunosuppressing Saccharibacteria to inflamed sites within the mucosal barrier, such as in gingivitis, to control inflammation and prevent further tissue damage^20^.

Oral bacteria deploy various strategies for surviving the oral mucosa, including immune suppression, pathogenic invasion, and evasion of immune killing through molecular masking and camouflage^33,38,46,129–131,152,153^. However, the combination of non-pathogenic endocytosis and suppression of proinflammatory pathways observed in TM7x has not been described before. TM7x uses a unique strategy to suppress immunoactivation by directly binding and inducing internalization of TLR2 receptors, and then downregulating TLR2 expression intracellularly. This strategy reduces available TLR2 on cell surfaces, preventing epithelial cells from mounting an effective response to other host or non-host bacteria. In combination with our previous studies showing that TM7x reduces production of antigenic molecules by their host-bacteria^20^, we begin to understand the potent ways in which Saccharibacteria modulate immune responses in the oral cavity. Perhaps this specialized role in mucosal immunology explains how Saccharibacteria have evolved alongside hominids throughout our entire evolutionary history, and why Saccharibacteria occur in oral cavities from many distantly related mammals^154,8,155^. Future studies will investigate if suppression of epithelial immunity by Saccharibacteria facilitates microbe-host immune balance therefore prevent disease, or facilitates assembly of dysbiotic microbiomes that contain oral pathogens and pathobionts that contribute to inflammatory diseases like periodontitis. Furthermore, downstream interaction with neutrophils and macrophages with Saccharibacteria still need to be investigated to fully understand the role of these bacteria in disease.

### Limitations of the study

For ethical reasons, potential causative agents of periodontitis cannot be tested on humans, so we utilized animal models and tissue culture studies. Currently, there are no well-established animal models specifically for oral mucosal immunity. To address this, our study uses multiple, independently derived human oral epithelial cell lines to test our findings. These cell cultures do not account for oral anatomy or include all the cell types present in human tissues. As such, these experiments do not reflect the natural aetiology of periodontal disease.

The human oral cavity harbours many of species of Saccharibacteria and Actinobacteria^8,43^ and we only tested the narrow subset of these strains that are cultivatable in the lab. We acknowledge that testing all Saccharibacteria species is impossible using cultivation-based methods. Future studies could address this by looking at oral microbiome communities.

## Supporting information

Suppmental figures 1-7

## Acknowledgements

This research was supported by grants from the <u>N</u>ational Institute of <u>D</u>ental and <u>C</u>raniofacial <u>R</u>esearch (NIDCR) under awards 1R01DE031274 (B.B.) and 1R01DE023810 (X.H., J.S.M.). Forsyth Institute Advanced Microscopy Core Facility supported by NIH 1S10OD034405-01 and Harvard Center for Nanoscale Systems (CNS) facilities. We acknowledge Dr. Ning Yu for providing the HGEPp cell line and Lujia Cen for general lab support. Jennifer Gundrum and Kyle Bredin provided technical support with microscopy and flow cytometry. Dr. Mary Ellen Davey and Dr. Hyun Young Kim provided technical support for using the NanoSight.

## Author contributions

Conceptualization by DC, XH, JSM, and BB. Data acquisition by DC, ASG, MK, AK, and LL. Methodology by DC, ASG, MK, KAK, KSS, PD, RJL, JSM, and BB. Formal analysis by DC, KAK KSS, PD, RJL, JSM, XH, and BB. Resources provided by RJL, JSM, XH, and BB. Data Curation by KAK, KSS, and JSM. Original Draft written by DC and BB. Review and Editing by DC, ASG, KSS, RJL, JSM, XH, and BB. Visualization by DC. Supervision by BB.

## Methods

### Lead Contact

Requests for further information, resources, and reagents should be directed to the corresponding author, Batbileg Bor.

### Material Availability

All unique reagents and bacterial strains generated in this study are available from the lead contact with completion of a Material Transfer Agreement.

### Data and code availability

The raw RNA sequencing data were deposited in GEO under the accession number GSE296366. Currently this data is not available to the public but will be available once the article is published.

## EXPERIMENTAL MODEL

### Bacterial culture

All bacterial strains, episymbiont containing cocultures, and growth conditions are listed in the key resources table. Briefly, before each experiment bacteria from frozen stocks were passaged three times in brain heart infusion (BHI) broth under microaerophilic conditions (2% O₂, 5% CO₂, 93% N₂) in a Whitley workstation at 37°C to ensure homogeneity^66^. For epithelial cells infections, overnight bacterial cultures from passage 3-7 were used. Bacteria were harvested and washed with PBS twice, adjusted to OD_600_ = 0.6, and plated on BHI agar plates to calculate bacterial colony forming units (CFU). For Saccharibacteria quantification, see Saccharibacteria quantification, viability testing and Mito-Tracker labelling.

### Human cell cultures and bacterial infection

Oral epithelial keratinocyte cells (TIGK, NOK-SI, HGEPp, and OKF-6), HEK293-Null, and HEK293-TLR2 cells were grown according to manufacturer defined culture conditions (see key resources table). Briefly, keratinocytes cells were grown in keratinocyte serum-free medium (K- SFM, Invitrogen, Carlsbad, CA), supplemented with 0.4 mM calcium chloride, 25 µg/mL bovine pituitary extract (BPE), 0.2 ng/mL epidermal growth factor (EGF), and a PSG antibiotic cocktail (Penicillin-Streptomycin-Glutamine). HEK293-Null and HEK293-TLR2 cells were grown in Dulbecco’s Modified Eagle Medium (DMEM) supplemented with 10% fetal bovine serum (FBS), 50 U/ml penicillin, 50 μg/ml streptomycin, 100 μg/ml normocin, and 10 μg/ml blasticidin. Cell cultures were incubated at 37°C in 5% CO₂. After each passage, cells were examined using phase contrast microscopy for cell density, morphology, and competency. Epithelial cells were seeded at a density of 0.1 X 10^6^ cells and at confluency cell number was ∼0.5 X 10^6^ cells in 12 well plate. For cell culture infections, isolated and cleaned XH001 (*S. odontolytica*), TM7x (*N. lyticus*), or coculture (XH001+TM7x) bacteria were added to cell cultures (see TM7x cleanup procedure). Unless specified, XH001 was added at a multiplicity of infection (MOI) of 10 (10:1; XH001:TIGK) and TM7x was added at an MOI of 50 (50:1; TM7x:TIGK).

To spatially separate cell cultures from bacterial cultures, two chamber infection vessels separated by a polyethylene terephthalate (PET) membrane with a 0.22 μm pore size were created. XH001 and coculture cells were introduced into an upper chamber, while confluent TIGK cells remained in the lower chamber, preventing cell-cell contact. Infections were incubated for 8 hours at 37°C in 5% CO_2_ prior to supernatant collection for cytokine quantification via ELISA assay (see Cytokine protein analysis). For immunomodulatory priming assays, epithelial cells were infected with a priming bacteria (TM7x or Actinobacteria controls) two hours before subsequent infection by a host or non-host Actinobacteria species. Conversely, for coinfection treatments, Actinobacteria and episymbionts were added simultaneously.

## METHOD DETAILS

### TM7x cleanup procedure

TM7x strains were isolated from cocultures as described previously^65,66,89^ with modifications to reduce/remove host bacteria debris contamination. Briefly, genetically homogenous host-episymbiont cocultures were inoculated into 200 mL BHI medium for overnight growth, pelleted at 3,000 x g for 5 minutes, and then supernatant was passed through a 0.45 µm PVDF filter (Stericup, Millipore, Cat #SCHVU01RE). Resulting filtrate was centrifuged at 20,000 x g for 1 hour. This centrifugation speed was chosen after technical optimization (20,000-80,000 x g, Figure S1) because it pelleted TM7x but removed most host bacteria material. Isolated TM7x were dialyzed in buffer (PBS) using a 1,000 kDa cut-off dialysis membrane to remove remaining BHI medium and host-derived impurities, which are themselves highly immunogenic (Figure S1B). This new protocol effectively reduces the abundance of contaminating immunogens present in cleaned TM7x, preventing non-specific immune activation (Figure S1B). Cleaned TM7x was stored at -80°C in a custom-designed freezing medium (5% calf serum + 5% glycerol in PBS) at an OD_600_ of 0.4 to maintain maximum viability.

### Saccharibacteria quantification, viability testing and Mito-Tracker labelling

To standardize MOIs, host and non-host Actinobacteria were plated on BHI agar plates to generate OD_600_ to CFU standard curves. Actinobacteria were also plated for all infection assays to ensure accurate counts. Saccharibacteria strains do not form colonies on agar plates; thus, culture independent quantification was developed. Isolated Saccharibacteria were diluted to OD_600_ 0.4, then quantified via qPCR with TM7 strain-specific primers^66^. In parallel, cleaned TM7x were also quantified using a NanoSight Pro (Malvern Panalytical, Model no. HBG5000). Standard curves were generated for relating OD_600_ to NanoSight detected particles and for relating OD_600_ to qPCR CT values (both curves shown in figure S1D). Saccharibacteria viability and cell integrity were assayed using 1 μM Sytox Green staining for 30 min at 37°C (Thermo Fisher Cat#S7020). Stained cells were washed with PBS, counter stained with DAPI, washed again, and observed for fluorescence on a Axio widefield microscope (Zeiss).

For fluorescent labelling using Mito Tracker Green or Mito Tracker Red (Invitrogen, Cat# M22425) bacterial cells (∼1*10^9^ cells/mL) were washed twice with PBS, suspended in BHI containing 100 nM MitoTracker dye, and incubated for 1 hour microaerophilically at 37°C. After MitoTracker labelling, bacteria were washed three times with PBS and visualized using the Axio Axio widefield microscope to ensure saturation of the bacterial population.

### Neutralization of bacteria via physical, chemical, and biochemical treatment

Actinobacteria and Saccharibacteria were physically neutralized via sonication (50% amplitude, 15 seconds ON 30 seconds OFF, 10 min) and via sample heating (95°C for 15 minutes). Actinobacteria and Saccharibacteria were chemically neutralized via two methods, 4% paraformaldehyde fixation and 100% ethanol fixation, both incubated for 30 minutes at room temperature. To enzymatically digest surface proteins, cells were treated with 50 μg/ml proteinase K for 30 min at 37°C, then protease activity was quenched by adding 1X broad spectrum serine, cysteine and metalloprotease inhibitor. To enzymatically digest bacterial surface polysaccharides, cells were treated with 100 μg/ml lysozyme and 25 U/ml mutanolysin for 30 minutes at 37°C. After enzymatic treatment, cells were washed and resuspended in PBS for viability analysis and infections. Treated bacteria were visualized via phase contrast microscopy (Nikon Eclipse E400) to confirm cell integrity and TM7x viability was quantified via Sytox green live-dead staining.

### Extraction of *S. odontolytica* lipoprotein

Actinobacterial lipophilic fractions were extracted using a previously developed TX-114 (Cat# 9036-19-5) extraction methodology^51^. Briefly, XH001 cells were cultured in BHI, washed in PBS, and resuspended in lipoprotein extraction buffer (150 mM NaCl and 10 mM Tris–HCl, pH 8.0). Resuspended cells were supplemented with 1/10 volume of 20% (vol/vol) aqueous TX-114, rotated at 4°C for 2 hours, and then centrifuged (15,000 × g, 5 min, 4°C) to remove cells. The supernatant was incubated at 37°C for 5 minutes and then centrifuged again to separate the lower (lipophilic) phase from the upper (aqueous) phase. Excess methanol was added to the lower phase to precipitate the lipophilic fraction. Precipitated mixtures were incubated overnight at −80°C, centrifuged to pellet precipitates (15,000 × g, 30 min, 4°C), decanted to remove supernatants, and resuspended in PBS (hereafter Lipo-fract). Lipo-fract protein concentration measured by Bio-Rad Protein Assay (Bio-Rad Laboratories, Hercules, CA).

### TLR2 agonist and antagonist treatment

Cultured TIGK and HEK293-TLR2 cells were dissociated into single-cell suspensions and cell viability (>83%) was analyzed using Trypan blue (0.4% solution) staining and hemocytometer-counting. Cells were seeded at the density of 0.1 X 10^6^ cells and at confluency the cell number was ∼0.5 X 10^6^ cells/mL in 12 well plate. Upon confluency, cells were treated with α-TLR2 antibody to neutralize TLR2 present on the cell surface. Confluent cells were washed with PBS and culture media containing 0.5 μg/ml α-TLR2 antibody was added and incubated at 37°C with 5% CO_2_ for 1 hour. After α-TLR2 antibody treatment, cells were washed with PBS then infected with XH001 (MOI 10), host-episymbiont coculture (MOI 10), or TM7x (MOI 50) (MOI is defined as # of infectious agents/# of compatible hosts). Infected cell cultures were incubated at 37°C with 5% CO_2_ for 8 hours, then supernatant was collected for cytokine protein detection (See Cytokine protein analysis). For TLR2 agonist treatment, confluent cells were treated with 20 μg/ml of pam2csk4 (TLR2/TLR6 agonist) or pam3csk4 (TLR2/TLR1 agonist) 37°C with 5% CO_2_ for 8 hours. For priming experiments, confluent cells were infected with TM7x for 2 hours at 37°C prior to TLR2 agonist treatment (pam2csk4 and pam3csk4) and standard incubation.

### Cytokine protein analysis

After each infection experiment, supernatants and cell lysates were collected to quantify cytokine protein and RNA transcript levels, respectively. For global cytokine protein analysis of human cytokine array, C5 kits were used according to manufacturer’s instructions (RayBio, see Key Resources table). Briefly, antibody array membranes were treated with blocking buffer for 1 hour (4°C), exposed to cell culture supernatants for two hours (room temperature), washed thrice, treated with biotinylated primary antibodies for two hours (room temperature), washed again, treated with HRP-streptavidin for two hours (room temperature), transferred to a plastic sheet, treated with detection buffer for two minutes (room temperature), and imaged via chemiluminescence. Densitometry analysis of expressed cytokines was performed using ImageJ software. Background signal was subtracted and data was normalized to the provided positive control. Heatmap analyzed and plotted using GraphPad Prism software version 10.4.2 (534).

Following global analysis, highly expressed individual cytokines (IL-8, GRO-α, MCP-1, IL-6, and TIMP2) from infected oral epithelial cells were measured using a human ELISA kit (R&D systems), as directed by the manufacturer. Briefly, wells in a 96 well plate were coated with captured antibodies overnight, washed thrice with wash buffer, incubated with blocking reagent for one hour (room temperature), exposed to 100 μl of supernatant collected from infected cell cultures in triplicate, and incubated for two hours (room temperature). Plates were treated with detection antibody, incubated for two hours (room temperature), treated with HRP-streptavidin, incubated for 30 minutes (room temperature), treated with TMB substrate, incubated for 15 minutes (room temperature), quenched with the stop solution, and quantified via 450 nm. Absorbance was normalized to the blank control and analysis was performed across a standard curve. Data was plotted using GraphPad Prism software version 10.4.2 (534).

### Cytokine transcript level analysis

Non-infected TIGK cell cultures were dissociated with trypsin into single-cell suspensions and cell viability (>83%) was analyzed using Trypan blue (0.4% solution) staining and hemocytometer counting. Cells were seeded for infection in 12 well plates. Upon confluency, cells were infected with XH001, TM7x, or coculture and incubated at 37°C, 5% CO_2_ for 8 hours. Total RNA was isolated from the infected cells using the Total RNA miniprep kit (NEB#T2110) as per manufacturer instruction. Briefly, treated cells were lysed using lysis buffer, RNA was collected in an RNA binding column then eluted. RNA qualities were assessed using Qubit (Invitrogen) and Nanodrop (ThermoFisher). Isolated RNA was stored at -80°C until use. cDNA was synthetized using the Invitrogen™ SuperScript™ II Reverse Transcriptase (Life Technologies) based on random hexamers, according to the manufacturer’s protocol and PrimeScript 1st Strand cDNA Synthesis Kit (Takara Bio) following the manufacturer’s protocol. The resulting cDNA was stored at -20°C for qPCR.

To quantify cytokine mRNA levels for induced by bacterial infection, the QuantStudio 3 and 5 Real-Time PCR systems (Applied Biosystems) were used to perform qPCR. GAPDH served as an internal control, and relative gene expression levels were calculated using the 2−ΔΔCt formula (ΔCT = Target Gene – Reference Gene)^156^. The following primers were utilized for the amplification of target genes: IL-8 (CXCL8): Forward primer: 5’-TGCCAAGGAGTGCTAAAGAAC-3’, Reverse primer: 5’-TCCACTCTCAATCACTCTCAGT-3’, Gro-α (CXCL1): Forward primer: 5’-GTCCGTGGCCACTGAACT-3’, Reverse primer: 5’- ATGACTTCGGTTTGGGCG-3’, MCP-1 (CCL2): Forward primer: 5’- CAATCAATGCCCCAGTCACC-3’, Reverse primer: 5’-GGGACACTTGCTGCTGGT-3’, CD14: Forward primer: 5’-CCACAGGACTTGCACTTTCC-3’, Reverse primer: 5’- CAGGTCTAGGCTGGTAAGGG-3’ and TNFAIP3: Forward primer: 5’- ACTCCCAAAGCTGAACTCCA-3’, Reverse primer: 5’-ACTTCATGGCAGTGGTCTCA-3’.

### Library preparation and total RNA sequencing

RNA isolations for meta transcriptomics were conducted using the Total RNA miniprep kit (NEB#T2110) as per manufacturer instruction and a brief protocol described above for RNA quantification. Subsequently, total RNA was prepared for sequencing. The samples were processed with the Microbiome Metatranscriptomics Sequencing Service (Zymo Research, Irvine, CA). The RNA-Seq library was prepared using the Zymo-Seq RiboFree, Total RNA Library Kit (R3000, Zymo Research, Irvine, CA) with 500 ng RNA as input. All libraries were quantified with TapeStation (Agilent Technologies, Santa Clara, CA) and then pooled in equal abundance. The final pool was quantified using qPCR. The final library was sequenced (∼40M PE reads per sample) on the NovaSeq® (Illumina, San Diego, CA) platform. Raw RNA-seq reads were first subjected to quality trimming to remove low-quality bases and adapter sequences. The resulting high-quality reads were then aligned to the human reference genome (*Homo sapiens* GRCh38, repeat-masked primary assembly, file: Homo_sapiens.GRCh38.dna_rm.primary_assembly.fa, downloaded from Ensembl in September 2023) using the Geneious RNA Mapper algorithm. This alignment method allows for reads to span across intronic regions of annotated coding sequences (CDS), enabling accurate mapping of exon-exon junctions. Additionally, reads that mapped ambiguously to multiple locations in the genome were retained and counted as partial matches, ensuring a more comprehensive representation of gene expression, particularly for genes with paralogs or repetitive sequences.

### Bioinformatic and sequence analysis

Raw counts per gene were input into the *DESeq2* Bioconductor/R package^157^ using the recommended guidelines, including filtering out genes with total counts less than 10 across all samples. Data were expressed as Log2 fold change values and P-values were adjusted for multiple comparisons using the Benjamini-Hochberg procedure^158^ (Table S1). Log fold-change shrinkage was applied to the output via *apeglm*^159^ refine the ranking of genes. This output was used to generate volcano plots using the *EnhancedVolcano* Bioconductor/R package (https://github.com/kevinblighe/EnhancedVolcano), and for functional enrichment analysis through the STRING Database version 12.0^160^ (FDR< 0.05; Interaction confidence score > 0.4). For heatmap and principal component analysis (PCA), raw counts were made homoscedastic using a regularized logarithm transformation^1^. PCA plots were generated using *ggplot2*^161^ and *ggforce*^161^ packages for R. Gene count data were transformed into z-scores and input into *ComplexHeatmap* Bioconductor/R package^162^. The TLR2 network heatmap was generated via STRING to include 49 direct interactors and subletting the RNASeq dataset. Heatmaps related to endocytosis and vesicle formation were generated by sub-sampling corresponding GO-term Biological Process pathways. Visualization of common gene sets between treatments was performed using the *UpSetR* R package^163^.

### siRNA transfection and cytokine quantification

TIGK cells were seeded into a 6-well plate at low density (∼0.05 X 10^6^ cells/mL) and allowed to grow until they reached 40-60% confluency. Once the desired confluency was achieved, old media was discarded and cells were washed with phosphate-buffered saline (PBS). Fresh Keratinocyte Serum-Free Medium (KSFM) was then added to each well to prepare for transfection. For samples subject to gene knockdown siRNA transfection mix was prepared for 6 well plate as follows: A total of 300 nM of siRNA was added from a 100 µM stock and combined with 1.2 ml of 1X LipoJet buffer and supplemented with 24 µL of LipoJet transfection reagent (SignaGen laboratories, Cat# SL100468). The transfection mixture was gently mixed and allowed to incubate at room temperature for 10 minutes. Following incubation, 200 µL of transfection mix was added to each well. Cells were immediately incubated in a CO₂ incubator (37°C, and 5% CO₂) for 48 hours. Post siRNA transfection, TIGK cells were washed with PBS and supplemented with fresh KSFM media prior to bacterial infection. XH001, TM7x, and coculture bacterial cells were added to each well at the appropriate MOIs (see TLR2 agonist and antagonist treatment). Pam2csk4 and Pam3csk4 were added to wells at final concentrations of 200 nM to provide positive controls.

### Flowcytometry based binding assay

Bacterial (XH001, TM7x, and coculture) labelling was optimized using MitoTracker Green/ Red as described above. To quantify oral epithelial cells associated with bacteria, flow cytometry-based analysis was performed. Oral epithelial cells were seeded in 6 well culture dishes, grown to confluence (1 × 10^6^ cells/well), washed with PBS, and infected with MitoTracker-labelled bacteria. At various time points post-infection, cells were detached via one minute accutase treatment (Cat# AT104). Detached cells were washed two times with ice-cold PBS and resuspended in 1 mL PBS. Samples were analyzed on a FACS Attune (Becton Dickinson) by gating on uninfected eukaryotic cells based on forward and side scatter. Cell-associated fluorescence was measured in fluorescence channel 1 (BL-1A and BL-1H) for 50,000 events/particles per sample, detecting MitoTracker-labelled cells. Signal data was recorded and analyzed using FlowJo software version 10.10. Side scatter was plotted on the y-axis and fluorescence plotted on the x-axis. Plotting and statistical analysis performed via GraphPad Prism software version 10.4.2 (534).

### Saccharibacteria genetics

All TM7x type IV pili mutants were generated via sequential transformation strategy as part of our previous study^1^. Briefly, linear constructs for gene deletion via homologous recombination were constructed using a hygromycin B resistance cassette alongside promoter and terminator regions from elongation factor Tu (pTuf) in *S. epibionticum*. For gene knockout constructs, 300 bp homology arms were PCR amplified, and linear constructs were generated using NEB HiFi Assembly. This hygromycin B resistance cassette was also inserted into a neutral site (NS1) in the TM7x genome as a control for the transformation process. For transformations, TM7x-XH001 cocultures were inoculated with 1 μg of linear DNA construct, incubated, then passaged with hygromycin B supplementation multiple times to enrich for transformants. Isolated mutants were confirmed by whole genome sequencing using Oxford Nanopore Technology and then isolated using the Saccharibacteria isolation protocol described above.

### Confocal imaging and analysis

Gingival epithelial cells were seeded over the sterile cover slip in 6 well plate at 0.3 × 10^6^ density and allowed to reach confluency. Confluent cells were washed and treated with fresh infection medium (culture medium without antibiotics) and TM7x (MitoTracker labelled/non-labelled) were introduced to TIGK cells and incubated for 2 or 8 hours before downstream treatments (see below). For experiments that used endocytosis inhibitors, chloroquinone, Dyngo- 4a, nystatin, and sodium azide were used at concentration of 50 µM, 5 µM, 20 µg/ml, and 100 µM prior to infection. To determine colocalization of endocytosis markers and TM7x, infected TIGK cells were fixed using 2% paraformaldehyde (PFA) to preserve cellular structures. Post fixation, cells were permeabilized by 0.5% Triton X-100 in PBS at room temperature for 10 minutes, followed by blocking with PBS containing 0.1% Triton X-100, 1% bovine serum albumin (BSA), and 10% donkey serum at room temperature for one hour. Subsequently, primary antibodies against EAA1, TLR2, LAMP-1, or LC3B were added at ratios of 1/2000, 1/500, 1/500, 1/500, and 1/500, respectively. Samples were incubated up to overnight. For colocalization imaging, after three washes (PBS supplemented with 0.025% Triton X-100), samples were incubated at room temperature for one hour with different color secondary antibodies: α-mouse IgG coupled to NL637 and NL437, and α-rabbit IgG coupled to Alexa Fluor 568 and CY3, all diluted 1/1000 in PBS containing 1% BSA and 0.1% Triton X-100. Cells are washed thrice after antibody treatment, incubated with Phalloidin diluted 1:500 with PBS for 15 minutes, incubated with DAPI diluted 1/2500 in PBS for 5 minutes, rinsed again, and suspended in ProLong™ Gold Antifade Mountant for microscopy. Confocal images were acquired using a Zeiss LSM 880 confocal laser-scanning microscope equipped with a super resolution 32-channel AiryScan detector. Samples were imaged using 633, 561, 488, and 405 nm excitation wavelengths at 63x magnification. To acquire z-stack images, z-step size was set at 180 nm and 10-30 slices were captured, covering ∼1.8-5.4 μm. Acquired images were processed with Zen Blue software and Image5D plugins in ImageJ software^164^. Maximum intensity projection of z-stack images were generated using false color unmixed channels for overlay images. 3D images were created on Zeiss LSM 980 microscope.

Briefly, z-stacked images were segmented, followed by surface and volume rendering to generate 3D models. Rendering parameters were adjusted for opacity, color, and lighting. daime analysis was done on TLR2, EAA1, and LAMP-1 immune stained TM7x infected groups. daime quantifies the spatial relationship of two populations by calculating a pairwise correlation value as a function of the distance between individual objects. Correlation values larger than 1 indicate populations that cluster together, values less than 1 indicate populations that cluster separately, and values around 1 indicate random distributions. Replicated pairwise correlation values were then used to calculate 95% confidence intervals for statistical testing (CI). Pearson’s coefficient was used to quantify colocalization of TM7x with TLR2, EAA1, and LAMP-1 in z-stacked images using the ImageJ software plugin JACoP. Analyzed outputs were plotted in GraphPad Prism software version 10.4.2 (534).

### Differential immune-staining of intra- or extra-cellular TM7x

To differentiate intra- and extracellular TM7x in infected TIGK cells, immune staining and super resolution microscopy were performed. Cell infection with TM7x was performed as described above. Infected cells were fixed using 2% paraformaldehyde (PFA) to preserve cellular structures and stored at 4°C for downstream TM7x staining. To stain TM7x, antibodies were raised against whole formaldehyde fixed TM7x (antigen) in rabbits (ProSci incorporated, Poway, California, USA). Post immunization period serum was extracted and run through a Protein A column to isolate IgG present. Isolated antibodies were then eluted and concentrated. Direct ELISA assays using fixed TM7x cells were run using rabbit anti-serum, immunodepleted serum (protein A column flow-through), and purified IgG antibody at various dilutions. Purified antibodies against TM7x were used for immunostaining TM7x infected TIGK cells. Post fixation, cells were washed with wash buffer and blocked with PBS containing 0.1% Triton X-100, 1% bovine serum albumin (BSA), and 10% donkey serum at room temperature for 1 hour. TM7x primary antibody (1:1000 dilution) was added in PBS containing 1% BSA and 0.1% Triton X-100 and incubated at 4°C for 2 hours. After three washes with wash buffer, samples were incubated at room temperature for 1 hour with secondary antibodies: α-rabbit coupled to CY3, diluted 1/1000 in PBS containing 1% BSA and 0.1% Triton X-100. After this first round of staining, cells were permeabilized with 0.5% Triton X-100 in PBS at room temperature for 10 minutes, followed by blocking with PBS containing 0.1% Triton X-100, 1% bovine serum albumin (BSA), and 10% donkey serum at room temperature for 1 hour. Again, TM7x primary antibody (1:1000 dilution) was added in PBS containing 1% BSA and 0.1% Triton X-100 and incubated at 4°C for 2 hours. Cells were washed again with washing buffer, then incubated with secondary antibody (α-rabbit coupled to Alexa Fluor 568) at the dilution of 1:1000 in PBS containing 1% BSA and 0.1% Triton X-100 and incubated at 4°C for 1 hour. Cells are washed three times, incubated with Phalloidin diluted 1:500 with PBS for 15 minutes, followed by DAPI diluted 1/2500 in PBS for 5 minutes, and rinsed them in PBS. Finally, we mounted them on glass slides using SlowFade gold antifade mountant. Confocal images were acquired using a super resolution Zeiss LSM 880 confocal laser-scanning microscope equipped with a 32-channel AiryScan detector, samples were imaged using 633, 561, 488, and 405 nm excitation wavelengths at 63x magnification. For z-stack analysis, step size was set to 0.18 μm and 10 to 30 slices were captured, representing a depth of 1.8 μm to 5.4 μm. Images were processed with Zen Blue software and Image5D plugins in ImageJ software^164^. Maximum intensity projections of z-stack images were generated by overlaying false color unmixed channels.

### Transmission electron microscopy

Gingival epithelial cells were seeded at a density of 0.3 × 10^6^ density in 6 well plate and allowed to grow until they reached confluency. Cells were thoroughly washed to remove residual media and suspended in fresh infection media. TM7x were introduced to the TIGK cells and incubated for 2 hours or 8 hours, alongside an untreated control group for comparison. Following each infection period, cells were fixed (2.5% glutaraldehyde, overnight at 4°C) to preserve cellular structures. Additionally, samples were treated with 2% electron dense osmium tetroxide in phosphate buffer to enhance TEM contrast. Fixed cells were embedded in agar to provide support during epoxy embedding. Specimens were dehydrated via graded concentrations of ethanol (30%-100%) followed by treatment with propylene oxide to enhance infiltration. Samples were embedded in Spurr’s resin, which was chosen to optimize penetration and ultra-thin sectioning. Semi-thin sections were cut using a glass knife^165^ and ultra-thin sections were produced via diamond knife sectioning. Ultra-thin sections were placed on copper grids^165^ and impregnated with uranyl acetate and lead citrate to further improve sample contrast. Samples were imaged using a Hitachi 7800 transmission electron microscope at the Harvard Center for Nanoscale Systems (CNS) facility. Resulting images were analyzed using ImageJ software, focusing on interactions between TM7x and gingival epithelial cell structures.

### Antibiotic protection assay for intracellular TM7x

Before the antibiotic protection assay, antibiotics capable of consistently killing TM7x had to be validated and optimized, identifying gentamicin and neomycin as effective selective agents. MitoTracker labelled TM7x cells lost detectable host infectivity after treatment with 50 µg/ml gentamicin and 50 µg/ml neomycin for 1 hour. Infectivity of host bacteria was quantified by quantifying the percentage of XH001 host cells bound by TM7x in phase contrast microscopy^27^.

TIGK cells at confluency were infected with MitoTracker-TM7x at 37°C, MOI 50, 5% CO_2_ for 4 hours. Infected TIGK cells were washed with PBS to remove unattached TM7x. Cells were treated with 50 µg/ml of gentamicin and 50 µg/ml neomycin for 1 hour to kill extracellular TM7x. These antibiotics cannot penetrate through the mammalian plasma membrane so intracellular TM7x would be protected^166,167^. After antibiotic treatment, cells were washed with PBS and fresh culture media added. To evaluate intracellular TM7x, differential immune-staining of infected cells were done at various time point (0, 8, 24 and 48 hours) as described above. Viable intracellular TM7x cells were released from TIGK cells by lysing them at 0, 8, 24 and 48 hours using lysis buffer containing 0.1% triton X-100 in PBS for 10 min. Cell lysates were centrifuged at 300 x g to remove TIGK cell debris and the supernatant was collected and split into two tubes: 1) for qPCR based TM7x quantification, and 2) for infecting XH001. 48 hours after infecting XH001, cultures were visualized via fluorescence microscopy to quantify fluorescently labelled TM7x on XH001 cells or free-floating in the medium. qPCR was performed using TM7x specific primer as described above.

For TM7x intracellular survival between TIGK generations, TIGK cells were infected with unlabeled TM7x and the antibiotic protection assay was performed as described above. However, instead of lysing TIGK cells post-infection, cells were left to grow and divide, before being passaged at confluency. Cells were detached using accutase and seeded to plate at 1:20 dilution from the total cells. Cells were passaged this way three times. At each passage, differential immune staining was used to visualize intracellular vs extracellular TM7. TIGK cell lysates were added to XH001 to test intracellular TM7x cell viability as described above and qPCR was used to quantify genome copies as described above. Data were plotted using GraphPad Prism software version 10.4.2 (534).

### Statistical Analysis

All experiments were done a minimum of three biological replicates. Statistical comparisons of group means were performed using one-way analysis of variance (ANOVA) to assess differences among multiple groups to correct for the increased risk of Type I error due to multiple comparisons, Bonferroni post hoc correction was applied using GraphPad Prism software version 10.4.2 (534). Significance levels are indicated as follows: *P* < 0.05 (*)*, P < 0.01* (\*\**), P < 0.001* (***), *P* < 0.0001 (****), and "NS" denotes results that were not statistically significant.

