## Supplementary material for "Episymbiotic Saccharibacteria suppresses epithelial immunoactivation through Type IV pili and TLR2 dependent endocytosis": Suppmental figures 1-7

**Figure S1.** Related to Figure 1.

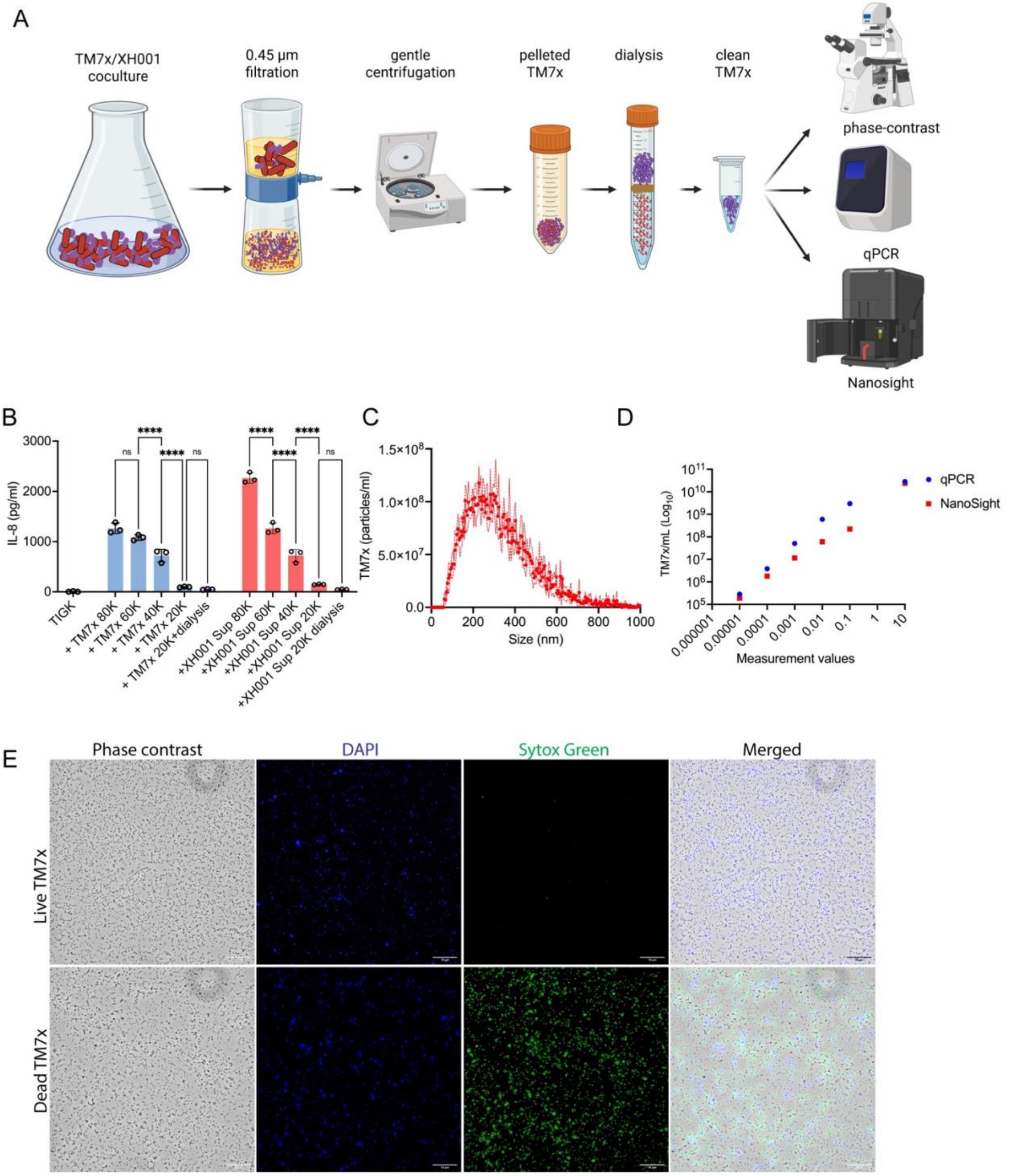


**Figure S1. Isolation of viable and clean Saccharibacteria.** (A) Graphical representation of our Saccharibacteria (TM7) isolation protocol for purifying episymbionts from host bacterium containing co-cultures. See Star methods for full detail. (B) Production of IL-8 by TIGK cells was quantified after treatment with isolated TM7x and XH001 supernatant (sup) to optimize the protocol for removal of antigenic contaminants. Various centrifuge speeds were tested to pellet TM7x without XH001 debris. (C) Biophysical characteristics of TM7x cells were measured using nanoparticle tracking analysis (Malvern Pananalytic NanoSight). TM7x cells ranged in size from 200 to 500 nm, with a mean diameter of ~300 nm. (D) We generated standard curves for relating optical density to cell density as reported by NanoSight quantification and cell density as reported by qPCR quantification. Frozen cells were stored at an $OD_{600}$ of 0.4, equivalent to 2.3 x $10^{10}$ cells/ml (NanoSight estimate) or 2.8 x $10^{10}$ cells/ml (qPCR estimate). (E) Live dead staining using Sytox green dye showed that ~99% of isolated TM7x were viable (see Star methods). Live and heat killed TM7x were stained with Sytox green (green) and DAPI (blue) before capturing fluorescent images. All experiments performed in biological triplicate. Means were compared via one-way ANOVA with Bonferroni correction for multiple comparisons with * $P<0.05$, ** $P<0.01$, *** $P<0.001$, **** $P<0.0001$, and NS = not significant. All scale bars are 10 $\mu$m.

**Figure S2.** Related to Figure 1.

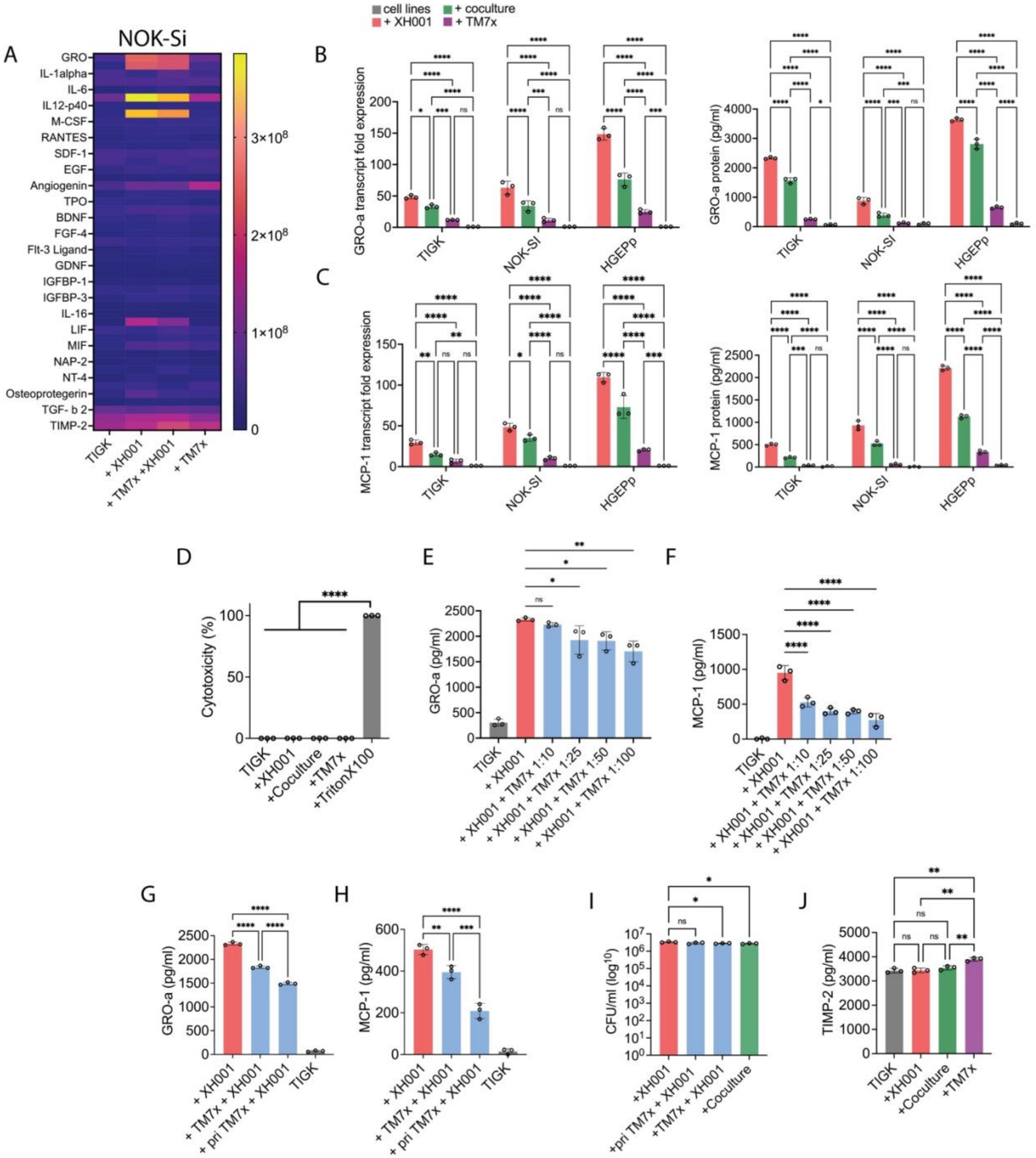


**Figure S2. Cytokine response and Saccharibacteria infection.** (A) Heatmap of global cytokine analysis of 81 human cytokine proteins for NOK-Si cells incubated with TM7x alone, XH001 alone, or TM7x/XH001 established coculture. The signal units are arbitrary gray value from the membrane array imaging (see methods). Focusing on cytokines Gro-$\alpha$ (B) and MCP-1 (C) demonstrates similar induction patterns in TIGK, NOK-SI, and HGEPp oral epithelial cells alone (gray), infected with XH001 (red), infected with TM7x (purple), or infected with the coculture (green). (D) Bacterial infection did not kill TIGK cells, as measured by lactate dehydrogenase (LDH) release, the Triton X-100 positive control killed ~100% of the cells. (E-F) TIGK cells (grey) infected with XH001 (red) or TM7x/XH001 co-cultures (blue bars; MOIs of 10, 25, and 50), then their production of (E) Gro-$\alpha$ and (F) MCP-1 were quantified. Production of (G) Gro-$\alpha$ and (H) MCP-1 were reduced more when TIGK cells were primed with TM7x prior to infection by XH001, relative to simultaneous infection (I) XH001 CFU count showing the bacterial load in TIGK cells infected with XH001 (red), primed TM7x with XH001 or simultaneous addition of TM7x and XH001 (blue) and coculture (green) groups post 8 hours of infection. (J) Production of TIMP-2 in TIGK cells alone (grey), infected with XH001 (red), infected with TM7x (purple), or infected with coculture (green). All experiments performed in biological triplicate. Means were compared via one-way ANOVA with Bonferroni correction for multiple comparisons with * $P<0.05$, ** $P<0.01$, *** $P<0.001$, **** $P<0.0001$, and NS = not significant.

**Figure S3.** Related to Figure 2.

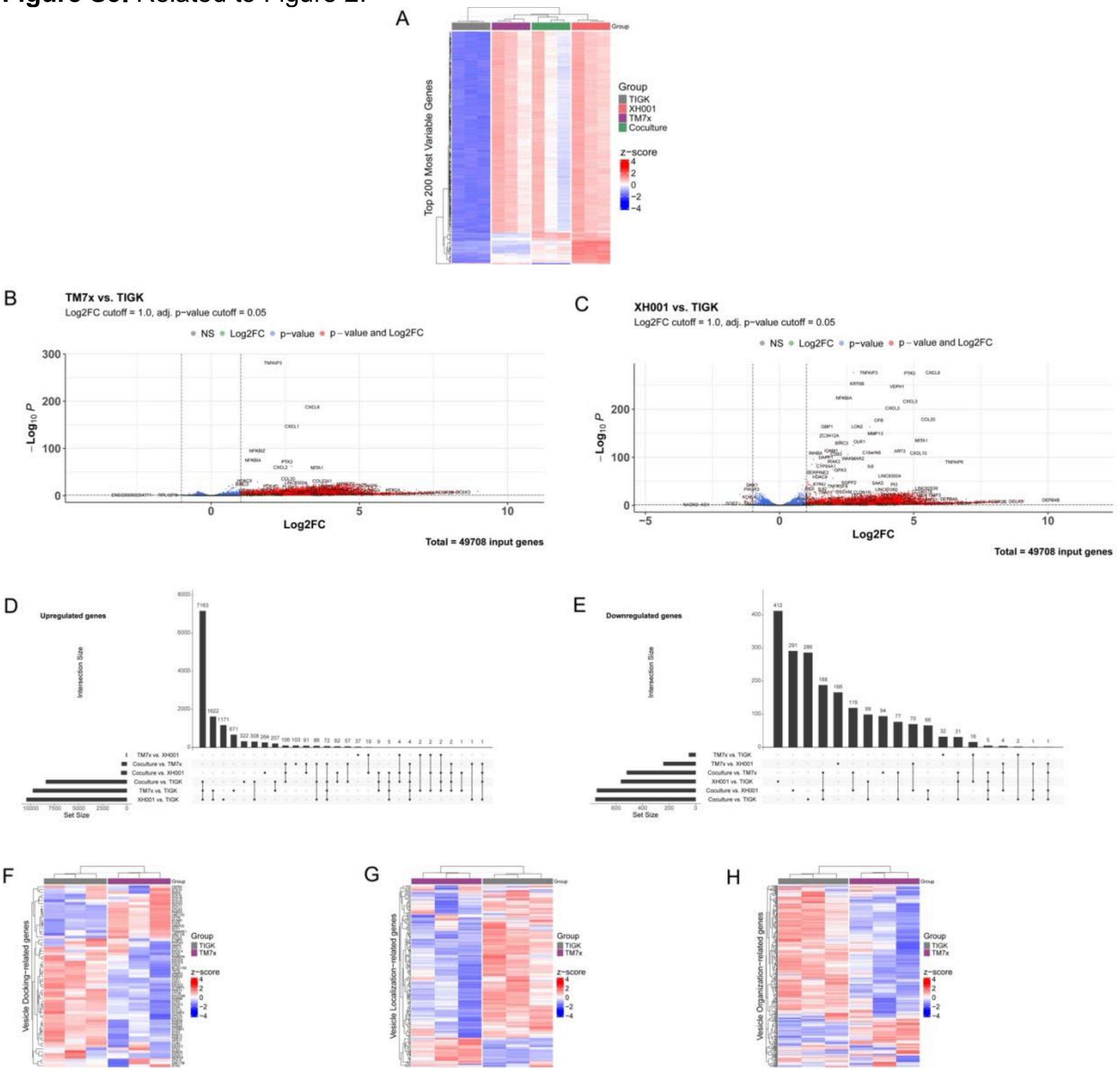


**Figure S3. Global gene response analysis of TIGK cells to XH001, TM7x and coculture.** (A) RNASeq data were normalized via log transformation and the 200 most differentially expressed genes were visualized via heatmap. Hierarchical clustering was performed to reveal similar groups according to both samples (columns) and genes (rows). XH001 infection induced elevated expression of more genes than infection by TM7x and coculture. In fact, many of the genes most strongly induced by XH001 infection were down regulated by TM7x infection. Volcano plot of differentially expressed genes after infection by (B) TM7x or (C) XH001 relative to uninfected TIGK cells. (D-E) Within each treatment, upregulated (D) and downregulated (E) genes were ranked and compared. Combining all three infection groups (XH001, TM7x, and the coculture)

reveals 7,163 upregulated genes unique to those treatments, here indicated with a dot. The downregulated genes, there is a dot on TM7x vs. Control and XH001 vs. Control, indicating that the corresponding bar in the bar graph shows 1,622 upregulated genes that are unique to those two comparisons that are not found in any other combination of comparisons. (F-H) RNASeq data were normalized via a $\log_{10}$ transformation, then vesicle-docking (F), vesicle localization (G), and vesicle organization (H) related genes were plotted on a heatmap for comparison. Hierarchical clustering was performed for columns (uninfected vs TM7x infected) and for rows (genes). Mirroring our principle component analysis, uninfected TIGK cell responses cluster separately from TM7x infected responses. Vesicle-mediated transports are perturbed by infection with TM7x and simultaneous infection by XH001 and TM7x induces a unique response.

**Figure S4**. Related to Figure 3.

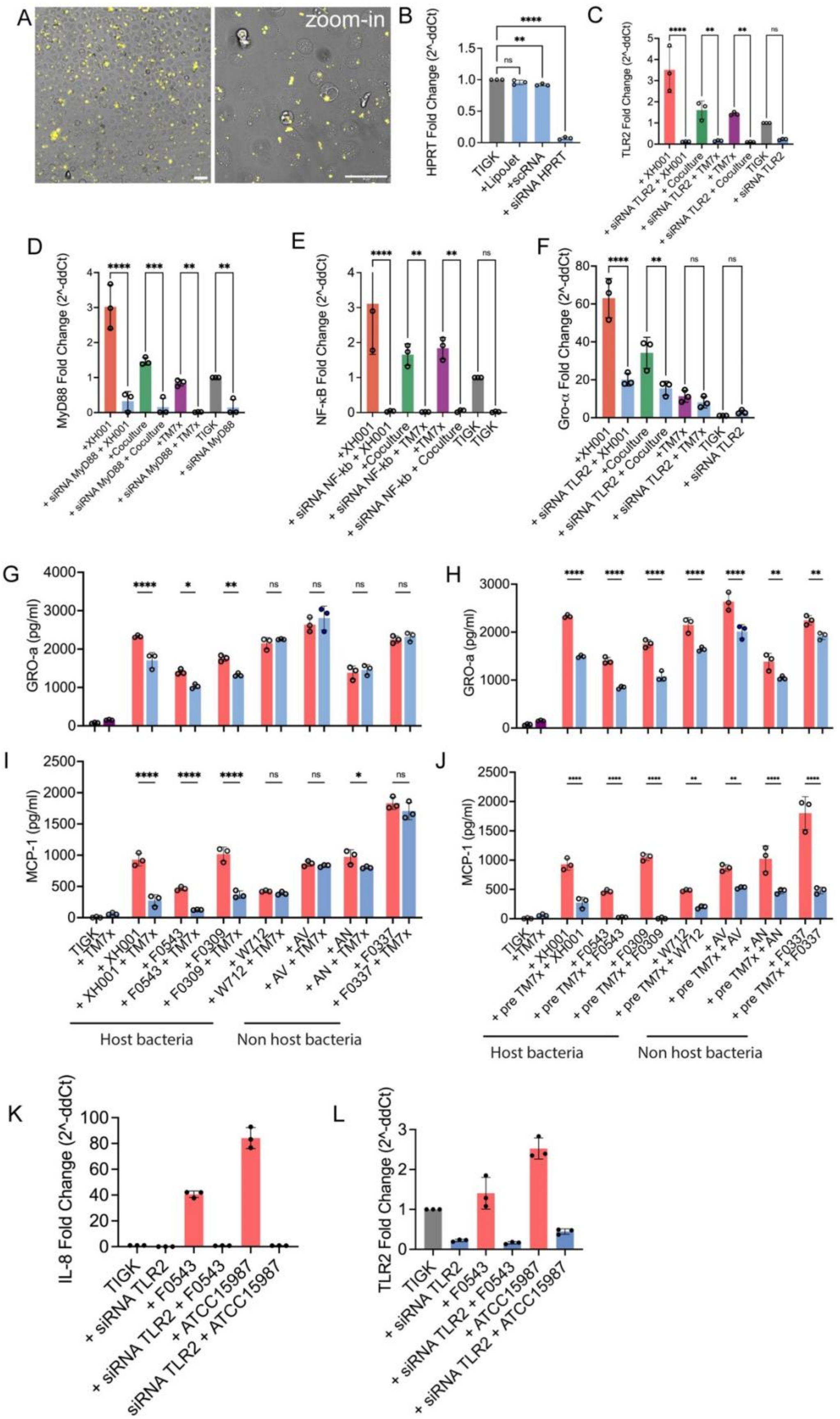

A
zoom-in
B
HPRT Fold Change (2^-ddCt)
TIGK
+LipoJet
+scRNA
+ siRNA HPRT
C
TLR2 Fold Change (2^-ddCt)
+ XH001
+ siRNA TLR2 + XH001
+ Coculture
+ siRNA TLR2 + TM7x
+ TM7x
+ siRNA TLR2 + Coculture
TIGK
+ siRNA TLR2
D
MyD88 Fold Change (2^-ddCt)
+XH001
+ siRNA MyD88 + XH001
+Coculture
+ siRNA MyD88 + Coculture
+TM7x
+ siRNA MyD88 + TM7x
TIGK
+ siRNA MyD88
E
NF-κB Fold Change (2^-ddCt)
+XH001
+ siRNA NF-kb + XH001
+Coculture
+ siRNA NF-kb + TM7x
+TM7x
+ siRNA NF-kb + Coculture
TIGK
TIGK
F
Gro-α Fold Change (2^-ddCt)
+XH001
+ siRNA TLR2 + XH001
+Coculture
+ siRNA TLR2 + Coculture
+TM7x
+ siRNA TLR2 + TM7x
TIGK
+ siRNA TLR2
G
GRO-a (pg/ml)
H
GRO-a (pg/ml)
I
MCP-1 (pg/ml)
TIGK
+ TM7x
+ XH001
+ XH001 + TM7x
+ F0543
+ F0543 + TM7x
+ F0309
+ F0309 + TM7x
+ W712
+ W712 + TM7x
+ AV
+ AV + TM7x
+ AN
+ AN + TM7x
+ F0337
+ F0337 + TM7x
Host bacteria
Non host bacteria
J
MCP-1 (pg/ml)
TIGK
+TM7x
+ XH001
+ pre TM7x + XH001
+ F0543
+ pre TM7x + F0543
+ F0309
+ pre TM7x + F0309
+ W712
+ pre TM7x + W712
+ AV
+ pre TM7x + AV
+ AN
+ pre TM7x + AN
+ F0337
+ pre TM7x + F0337
Host bacteria
Non host bacteria
K
IL-8 Fold Change (2^-ddCt)
TIGK
+ siRNA TLR2
+ F0543
+ siRNA TLR2 + F0543
+ ATCC15987
siRNA TLR2 + ATCC15987
L
TLR2 Fold Change (2^-ddCt)
TIGK
+ siRNA TLR2
+ F0543
+ siRNA TLR2 + F0543
+ ATCC15987
+ siRNA TLR2 + ATCC15987

**Figure S4**. **TLR2 signaling pathways are required for induction of cytokine response.** (A-B) The efficiency of LipoJet mediated siRNA transfection of TIGK cells was validated by (A) microscopy imaging of the fluorescently labelled siRNA (TYE 563) in siRNA transfected cells, and (B) transcriptome fold change analysis of HPRT gene in TIGK epithelial cells. (C-E) Knockdown of TLR2 (C), MyD88 (D) and NF-κb (E) in TIGK cells were conducted using siRNA. Expression of TLR2, MyD88, and NF-κb genes in siRNA transfected (blue) and non-siRNA transfected TIGK cells were quantified for uninfected cell cultures (grey), XH001 infected cells (red), TM7x infected cells (purple), and coculture infected cells (green). (F) Transcriptional expression of Gro-α in TLR2-siRNA transfected TIGK cells (blue) and non-siRNA transfected TIGK cells for uninfected (grey), XH001 infected (red), TM7x infected (purple), and coculture infected (green) cell cultures. Gro-α (G) and MCP-1(I) expression in TIGK cell responding to Actinobacteria infection with (blue) and without (red) simultaneous episymbiont infection. Priming TIGK cells with TM7x infection prior to addition of Actinobacteria reduced Gro-α (H) and MCP-1(J) expression more than the simultaneous infections. (K-L) TIGK cells siRNA transfected to knockdown TLR2 (blue) showed drastic reduction of TLR2 expression (K) and IL-8 expression (L) when uninfected (grey), infected with F0543 (red), or infected with ATCC15987 (red). All experiments performed in biological triplicate. Means were compared via one-way ANOVA with Bonferroni correction for multiple comparisons with * $P<0.05$, ** $P<0.01$, *** $P<0.001$, **** $P<0.0001$, and NS = not significant. All scale bars are 10 μm.

**Figure S5**. Related to Figure 4.

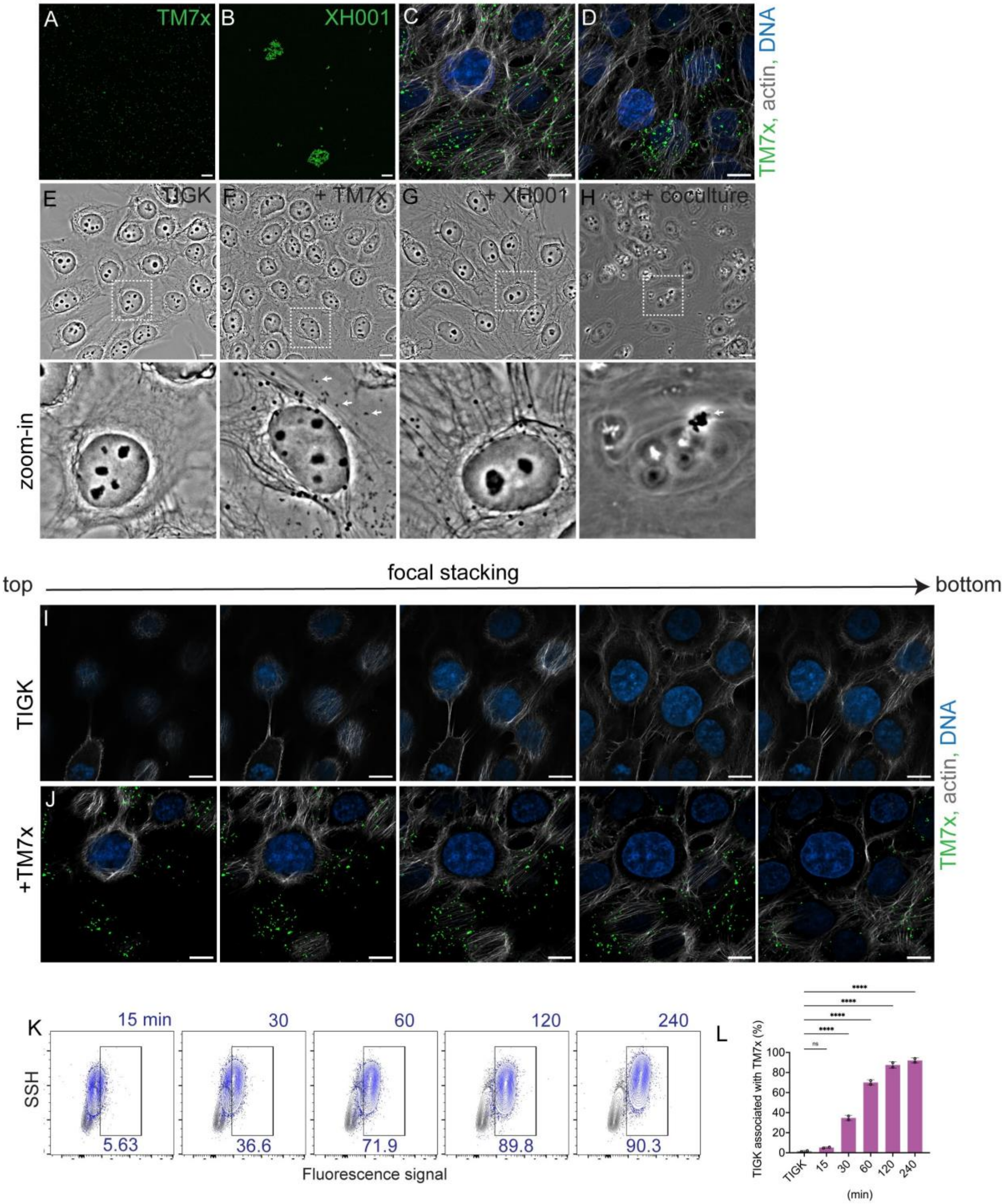


**Figure S5. TM7x exhibits strong adhesion to oral epithelial cells.** (A) TM7x cells and (B) XH001 cells were labelled with MitoTracker Green fluorescent dye to visualize epithelial association. (C-D) Addition of phalloidin to stain actin (grey) and DAPI to stain nuclei (blue) suggested that the cell associated TM7x could be intracellular. (E-H) Phase contrast images showing bacterial attachment to TIGK epithelial cells: (E) uninfected TIGK cells, (F) TM7x-infected, (G) XH001-infected, and (H) coculture infected treatments. White dashed boxes indicate the origin points for the zoomed in images presented below. Analysis of individual z-stack images from uninfected (I) and TM7x infected (J) TIGK cells also suggest intracellular localization. (K-L) Using flow cytometry to detect fluorescent TM7x at several time points post epithelial cell infection showed that epithelial binding starts rapidly, within 15 minutes, and takes approximately 2 hours to reach saturation. All experiments performed in biological triplicate. Means were compared via one-way ANOVA with Bonferroni correction for multiple comparisons with * $P<0.05$, ** $P<0.01$, *** $P<0.001$, **** $P<0.0001$, and NS = not significant. All scale bars are 10 μm.

**Figure S6**. Related to Figure 4

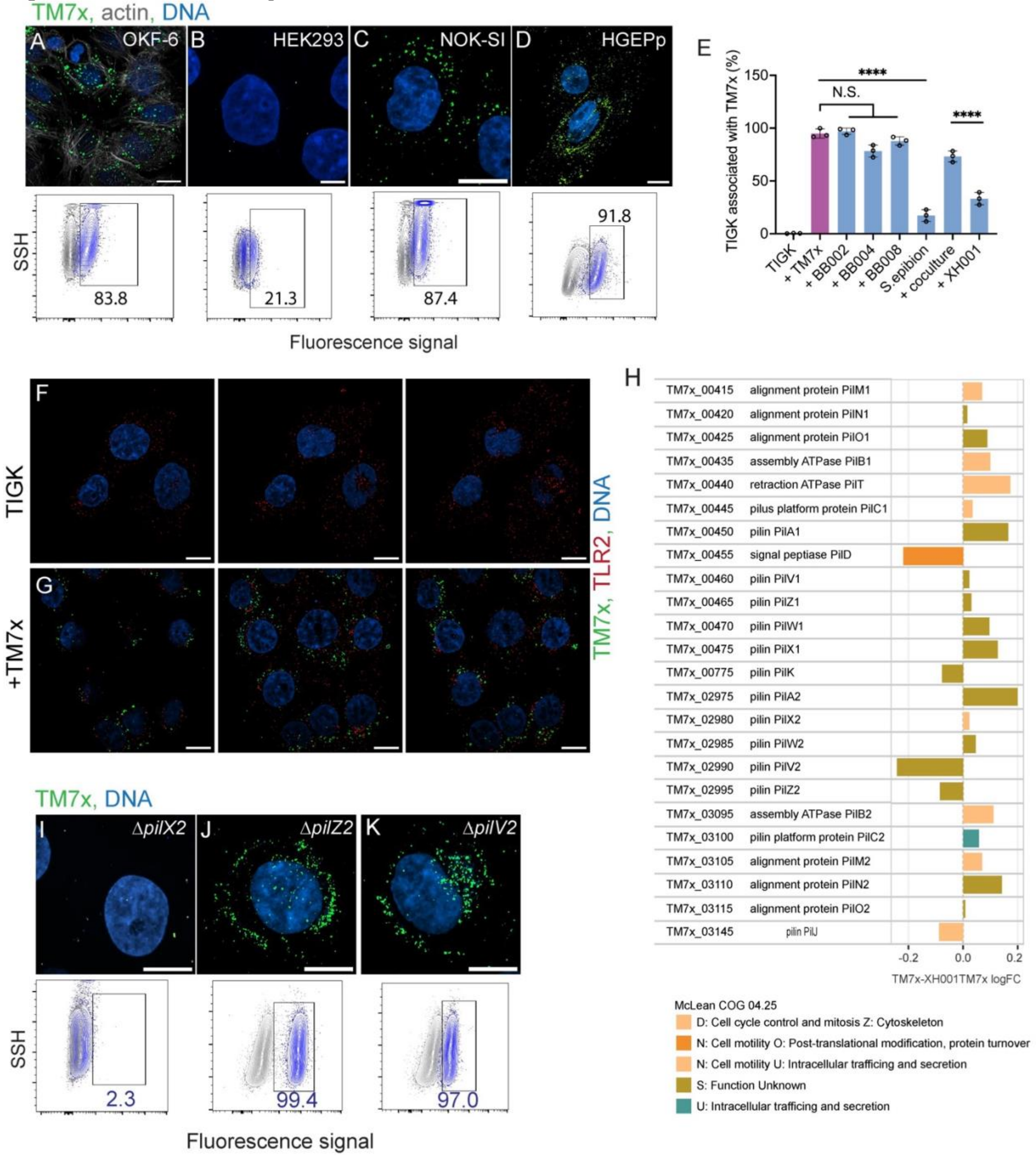


**Figure S6**. **TM7x type IV pili and TLR2 interaction with gingival epithelial cells.** (A-D) Fluorescence images and flow cytometry scatter plots depicting multiple epithelial cell lines infected with TM7x. All cell lines except HEK293 showed high levels of TM7x association. (E) Infecting TIGK cells with a range of oral Saccharibacteria (TM7x-purple, BB002-blue, BB004-blue, BB008-blue, *S. epibionticum*-blue) showed that most strains associated with epithelial cells, however *S. epibionticum* and host Actinobacteria showed very little association. (F-G) Z-stack images of TIGK cells alone (F) reveal a uniform distribution of TLR2 receptors (red) throughout cells. In contrast, TM7x-treated TIGK cells (G) show strong colocalization of TM7x (green) with the TLR2 receptor (red) across all Z-stack slices. (H) Comparison of TM7x T4P gene expressions from our transcriptomic analysis between groups TM7x alone versus TM7x-XH001 coculture with TIGK cells. Positive LogFC indicate genes that are expressed higher in TM7x alone group compared to coculture. D, N, S and U are functional COGs that these genes fall under. (I-K) Microscopy and flow cytometry analyses demonstrating that loss of type 4 pili genes disrupts epithelial cell binding byTM7x. All experiments performed in biological triplicate. Means were compared via one-way ANOVA with Bonferroni correction for multiple comparisons with * $P<0.05$, ** $P<0.01$, *** $P<0.001$, **** $P<0.0001$, and NS = not significant. All scale bars are 10 $\mu$m.

**Figure S7**. Related to Figure 5.

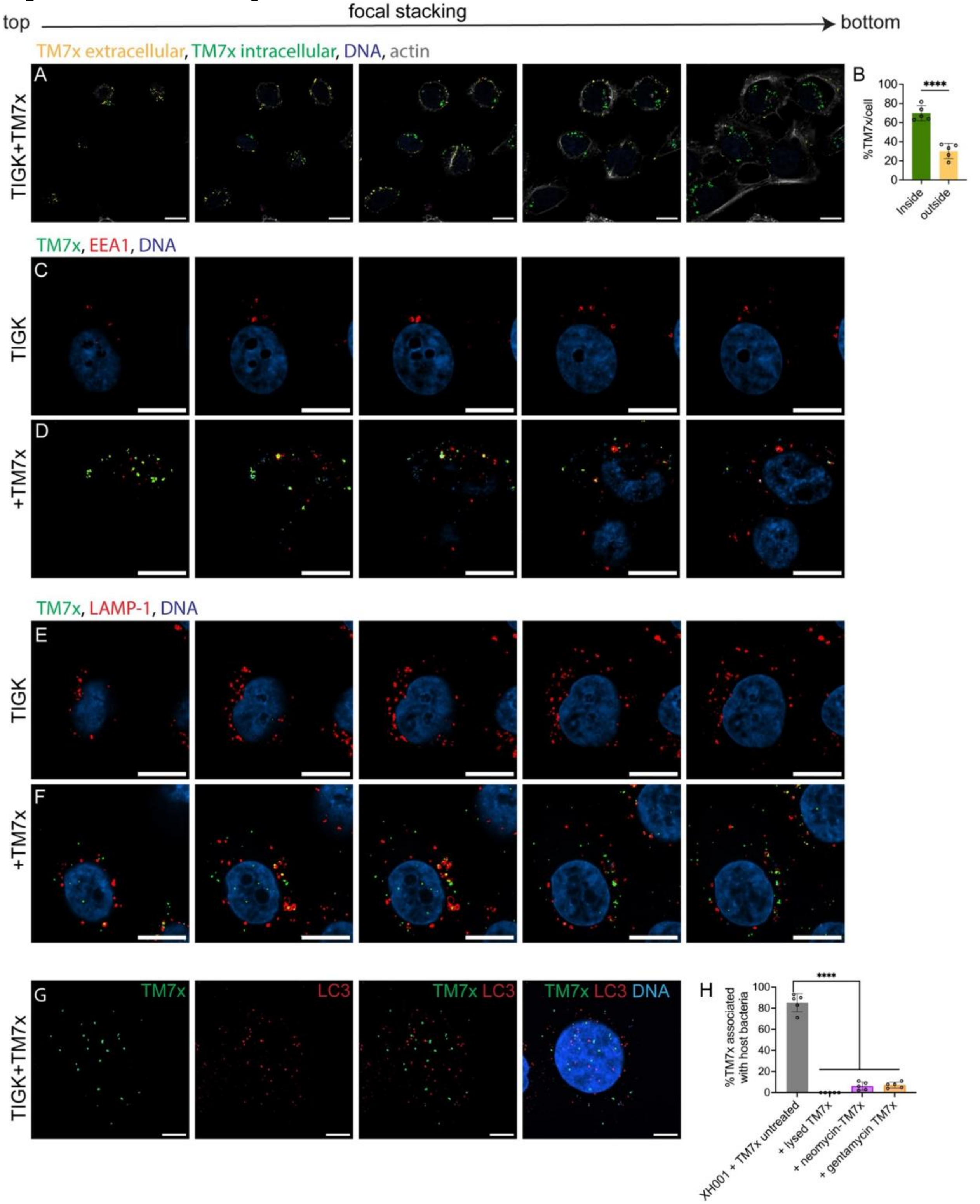


**Figure S7**. **TM7x internalized into epithelial cells via endocytosis.** (A-B) Differential staining of TM7x infected TIGK cells highlights the localization of TM7x bacteria as transition through the Z-stack. (A) After staining, extracellular TM7x appear yellow and intracellular TM7x are green, allowing clear segmentation of these populations and comparison of their positions relative to actin (grey) and DNA (blue). (B) Quantification of these populations showed ~60-70% of TM7x were intracellular. Z-stack analysis of TIGK cells immune-stained for the endosomal marker proteins EEA1 (C-D) and LAMP-1 (E-F) indicate endosomal localization of intracellular TM7x. (G) Autophagy marker protein LC3 (red) did not colocalize with TM7x (green). (H) Neutralization of TM7x via with neomycin treatment (purple), gentamycin treatment (yellow), and lysis (blue) all ablated epithelial cell binding/endocytosis relative to untreated TM7x (grey). All experiments performed in biological triplicate. Means were compared via one-way ANOVA with Bonferroni correction for multiple comparisons with * $P<0.05$, ** $P<0.01$, *** $P<0.001$, **** $P<0.0001$, and NS = not significant. All scale bars are 10 μm.
